# Development of tools to detect and identify strains belonging to the *Pseudomonas syringae* species complex responsible for vein clearing of zucchini

**DOI:** 10.1101/2023.05.02.539078

**Authors:** Caroline Lacault, Martial Briand, Marie-Agnès Jacques, Armelle Darrasse

## Abstract

Vein clearing of zucchini (VCZ) is a seed-borne bacterial disease that affects young plants of *Cucurbita pepo* subsp. *pepo*. VCZ agents are distributed into four phylogenetic clusters within the clades 2a and 2ba of the phylogroup 2 of *Pseudomonas syringae* species complex. Strains belonging to clades 2b and 2d are sometimes isolated from zucchini seeds but have not been associated with VCZ epidemics. Development of tools able to identify VCZ agents is important to better control the disease. Primers were designed to implement a seven-gene MLSA scheme on a collection of strains isolated from zucchini seeds. A clear predominance of strains with a host range on cucurbits limited to the genus *Cucurbita* (cluster 2ba-A) and the presence of VCZ strains in a fifth cluster (2ba-C) were evidenced. PCR tests were designed to characterize VCZ clusters and a multiplex qPCR test was proposed to distinguish strains having narrow cucurbit host range, associated to the presence of *avrRpt2* and *sylC,* from broad host range strains associated to the presence of *hopZ5* and *sylC*. Additional qPCR tests targeting clade 2b and 2d were also designed to gain insights on *P. syringae* strains that could be isolated from cucurbits. Specificity of these tools was evaluated *in silico* on the whole NCBI database and *in vitro* on a strain collection, showing a 100% inclusivity, except for the test dedicated to clade-2b strains, and an exclusivity ranging from to 96.7% to 100%. These different tools are intended to serve phylogenetic studies, epidemiological monitoring and seed testing.

## Introduction

Vein clearing (VCZ) is a bacterial disease that affects young plants of zucchini (*Cucurbita pepo* subsp. *pepo*), leading to plantlet deformations, water-soaked and necrotic spots on cotyledons and leaves, and stunting (Lacault et al. 2020). A transient symptom of vein clearing may be observed on the young first leaves but no symptoms have been so far reported on adult field-grown plants (Manceau et al. 2011; Lacault et al. 2020). VCZ is a seed-transmitted disease, as a clear link has been established between sowing infected seed lots and disease outbreaks in nurseries, and seed infection negatively impacts germination (Manceau et al. 2011).

The causal agents of VCZ belong to the *Pseudomonas syringae* species complex (Lacault et al. 2020) and are closely related to strains leading to symptoms on adult cucurbit plants such as systemic infection of zucchini (Djitro et al. 2022b) and bacterial leaf spots (Newberry et al. 2019; Qiao et al. 2022). *P. syringae* species complex (Gomila et al. 2017) includes bacteria that cause serious epidemics on woody plants and annual crops (for reviews Lamichhane et al. 2014, 2015) but also many strains that are present in the environment, some of which being phytopathogenic (Morris et al. 2013). Taxonomy of the *P. syringae* species complex has been revisited with the description of 13 phylogroups (PGs) subdivided in clades (Berge et al. 2014) and 19 phylogenomic species have been proposed (Gomila et al. 2017). Comparative genomics of strain collections has improved the knowledge of structure and evolution of this species complex highlighting impacts of recombination and horizontal gene transfer events. Homologous recombination blurs boundaries of some species (Dillon et al. 2019b). Based on former studies that identified and characterized different functions involved in pathogenicity (i.e. toxins, type three secretion systems (T3SSs) and repertoires of type three effectors (T3Es)), comparative genomics allows to discriminate bacteria with different behaviors such as specialized pathogens with a canonical T3SS, numerous T3Es and few toxins from more generalist ones with fewer T3Es but more toxins; some strains also lack a canonical T3SS and are mostly non-pathogenic (Baltrus et al. 2017; Dillon et al. 2019a; Xin et al. 2018).

VCZ strains are genetically diverse as the genome sequences of these strains are distributed in four clusters of the *P. syringae* species complex (Lacault et al. 2020). Clusters 2ba-A and 2ba-B belong to the clade 2ba (Lacault et al. 2020), which is issued from homologous recombination between clades 2a and 2b (Newberry et al. 2019). Clusters 2a-D and 2a-E belong to clade 2a (Lacault et al. 2020) corresponding to *P. cerasi* (Gomila et al. 2017). These clusters also encompass strains that are pathogenic on other cucurbits and share Average Nucleotidic Identities based on Blast (ANIb) values with VCZ strains as high as those observed between two VCZ strains (Lacault et al. 2020; Newberry et al. 2019). Two closely related clusters, clusters 2ba-C and 2a-F, group strains that have been sampled from watermelon, cantaloupe and squashes in the USA (Newberry et al. 2019). Recently, clusters 2ba-A and 2ba-C strains causing unusual symptoms on zucchini, such as twisted petioles, necrotic leaves, crown-rots and internal fruit rots have been reported in Australia (Djitro et al 2022a, b). An extended MultiLocus Sequence Analysis (MLSA) scheme has been proposed to allocate strains in these clusters and facilitate the analysis of large strain collections (Lacault et al. 2020). It includes to the previously available scheme based on four housekeeping genes (*gapA, gltA, gyrB*, and *rpoB,* Hwang et al. 2005), three largely distributed genes (*Psyr3420, Psyr4880* and *Psyr3208*).

The various VCZ clusters of strains have different host range extents. VCZ strains from cluster 2ba-A are characterized by a narrow host range of cucurbits that is limited to the genus *Cucurbita*. All other VCZ strains have a wider cucurbit host range, affecting squash (*Cucurbita* sp.) but also melons (*Cucumis mel*o), cucumbers (*Cucumis sativus*) and sometimes watermelons (*Citrullus lanatus*) (Lacault et al. 2020). The extent of these host ranges is consistent with their repertoire of T3Es. In particular, cluster 2ba-A strains, having the narrowest host range, are characterized by the presence of *avrRpt2*, while the clusters with a wider host range harbor *hopZ5*. Similarly, Djitro and colleagues (2022a) also report that Australian strains, harboring *avrRpt2* (from cluster 2ba-A and part of cluster 2ba-C, proposed to be named 2ba-C2), have a host range of cucurbits more limited than other 2ba-C strains harboring *hopZ5* (proposed to be named 2ba-C1).

Besides these main genetic clusters, a few strains isolated from zucchini seed lots are scattered in the clades 2b (Lacault et al 2020) and 2d (this study), corresponding to the *P. syringae* species *sensu stricto* and to the new genomic species A (Gomila et al. 2017), respectively. One isolated 2b strain causing symptoms on zucchini is genetically close to clade-2b strains belonging to the pathovars *aptata* and *syringae* (Lacault et al. 2020), which are reported to be pathogenic on cantaloupe and squashes (Morris et al. 2000; Langstone et al. 2003; Sedighian et al. 2014). Clade-2d strains isolated from watermelon and cantaloupe plants in the USA and France, are rather aggressive on their isolation hosts (Berge et al. 2014; Newberry et al. 2016), and in contrast weakly aggressive on squashes (Newberry et al. 2016). Seed testing is a first step to control seedborne bacterial diseases and requires specific and sensitive detection tools to detect pathogenic bacteria among a variety of microorganisms, including look-alikes or closely related strains that are not pathogenic, but are seedborne (Jacques et al. 2012). Currently, seed-associated plant pathogenic bacteria are mainly detected using PCR-based tests, due to their ease of use and specificity. A large range of PCR-based detection and identification tools targeting different phylogenetic groups of the *P. syringae* species complex have been developed (Guilbaud et al. 2016; Borschinger et al. 2016), as were PCR tests targeting genes encoding toxins and other pathogenicity factors (Bultreys and Gheysen 1999; Bereswill et al. 1994; Prosen et al. 1993; Chen et al. 2020; Galleli et al. 2014; Meng et al. 2014). A qPCR test targeting *sylC* (4Ba marker) was designed to detect VCZ strains (Manceau et al. 2011). Indeed, *sylC* is one of the genes encoding the non-ribosomal peptide synthetase (NRPS) responsible for syringolin synthesis (Amrein et al. 2004). The syringolin biosynthesis genes are widely distributed in phylogroup 2 strains (Baltrus et al. 2011; Dillon et al. 2019b; Dudnik and Dudler 2014; Hulin et al. 2018). This marker may hence not be specific enough for VCZ strain detection.

The aim of the study was to design and evaluate new identification and detection tools for VCZ strains. First, we developed PCR tests to amplify three genes to be included in the seven-genes MLSA scheme and characterized our collection of VCZ strains with this scheme. Second, we gained insight on clade-2d strains isolated from zucchini seeds, for which very little data were available, by sequencing two genomes. Third, faced to the genetic diversity of the strains responsible for VCZ, we used two approaches to design identification and detection tests. The first approach was based on the search for phylogenetic markers of the clusters and phylogenetic groups, which the VCZ belong to. The second one was to design a multiplex Taqman qPCR assay that associated to *sylC* other pathogenicity markers such as *avrRpt2* and *hopZ5,* whose presence characterized VCZ strains according to their host range on cucurbits. Finally, the multiplex qPCR test was evaluated for detection in seed samples.

## Materials and Methods

### Bacterial strains and growth

A bacterial collection of 112 strains including 54 strains responsible for VCZ (Lacault et al. 2020), 14 strains isolated from zucchini seeds or seed-production fields and belonging to the clade 2b (six strains) and clade 2d (eight strains), 35 strains from diverse *Pseudomonas* species and pathovars, seven bacterial strains pathogenic on cucurbits belonging to divergent genera (*Pectobacterium, Erwinia, Ralstonia, Xanthomonas,* and *Acidovorax*), and two strains associated with cucurbit seeds belonging to *Bacillus* and *Paenibacillus,* (Kalhaf et Raizada, 2016) was used in this study (Table 1). Representatives of other genus, species and pathovars were provided by the French Collection of Plant-Associated Bacteria (https://cirm-cfbp.fr/).

**Table 1.**
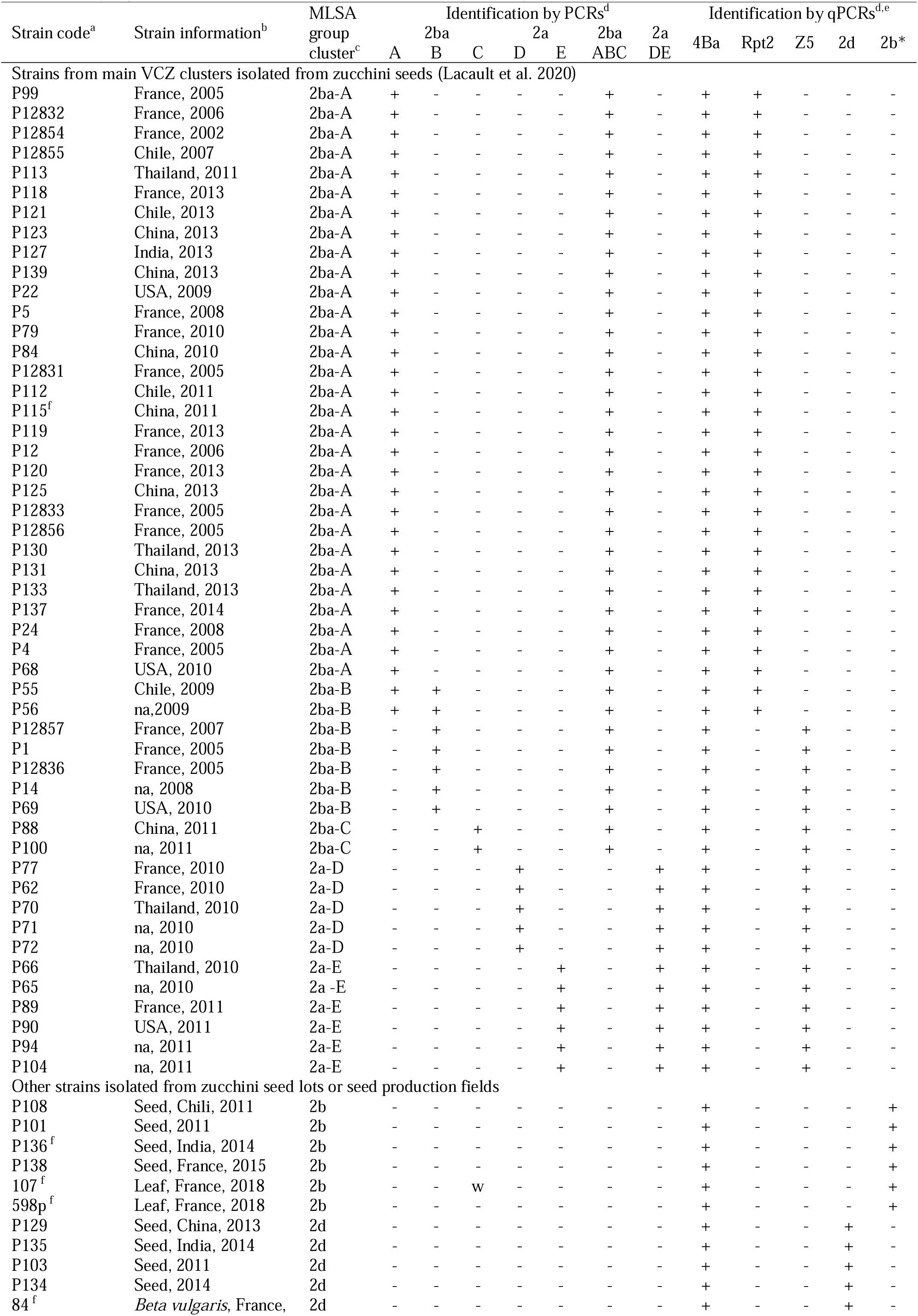
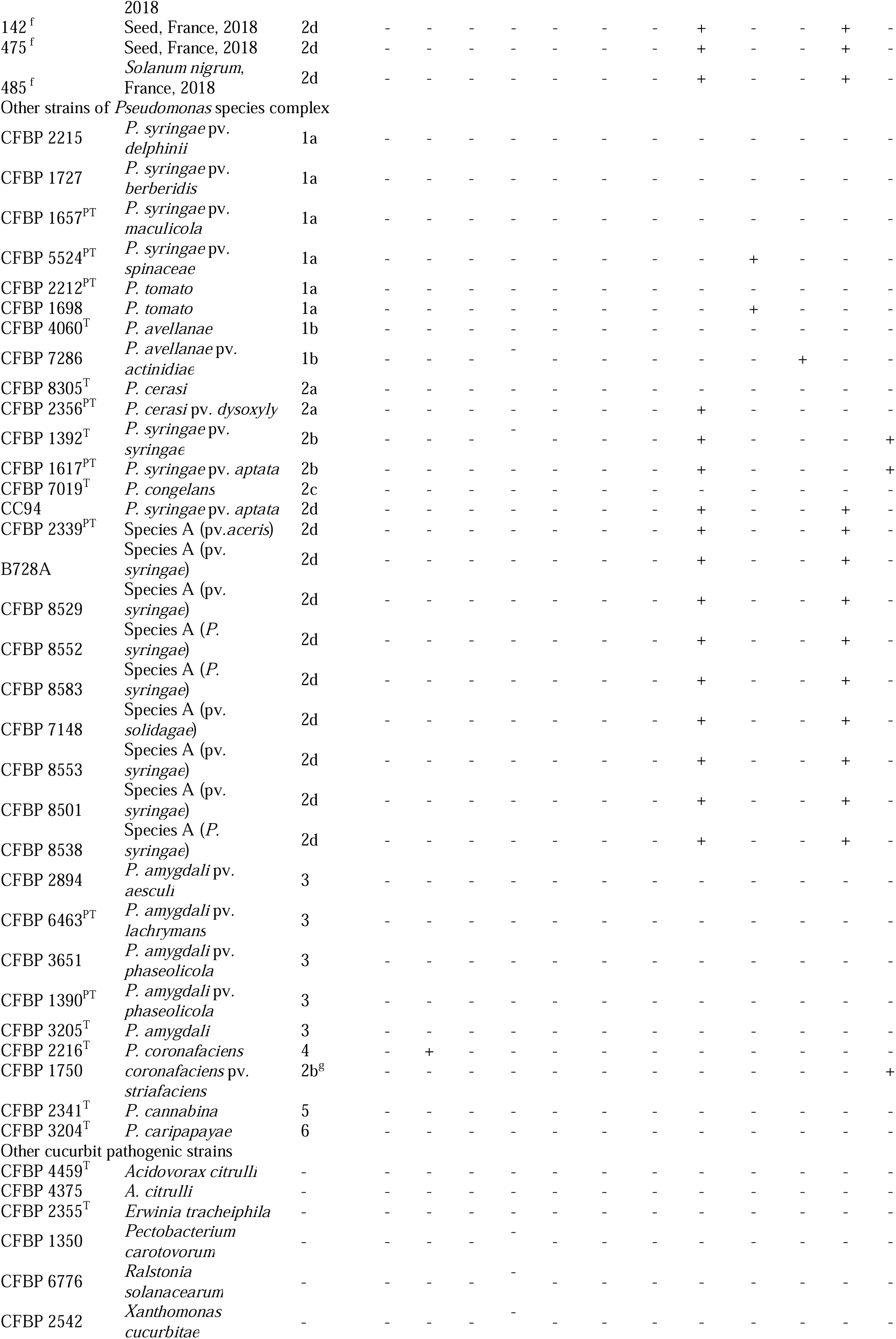
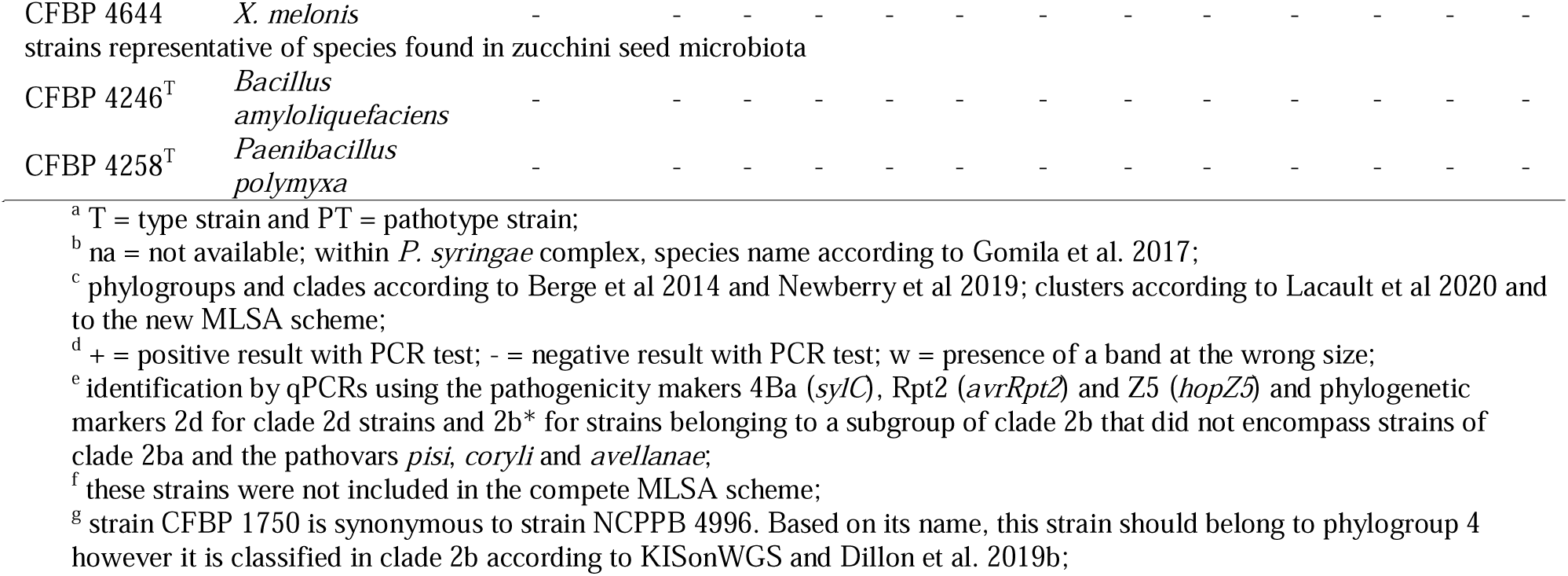
Phylogenetic position of bacterial strains used in this study and results of detection assays

Bacterial strains were stored in a −80°C freezer in 40% glycerol solution and were routinely cultivated at 28°C on 10% TSA medium (3 g.L^-1^ tryptone soya broth and 15 g.L^-1^ agar) with 50 mg.L^-1^ cycloheximide. Bacterial suspensions were prepared from fresh cultures suspended in sterile distilled water, calibrated at OD_650 nm_ = 0.1 (around 1 × 10^8^ CFU.mL^−1^), and adjusted to the final desired concentration with sterile distilled water. Dilution plating and colony counts after 48-h growth on 10% TSA were used to precisely quantify bacterial population sizes. Calibrated suspension-dilutions were boiled during 10 min and used as standards for qPCRs.

### Amplification and sequencing of the three genes to be included in the MLSA scheme

Three genes have been described for typing VCZ strains (Lacault et al. 2020). In the genome sequence of strain B728a, these genes correspond to (i) *Psyr3208* encoding the NADH dehydrogenase subunit M, (ii) *Psyr3420*, encoding a protein belonging to FixH superfamily, and (iii) *Psyr4880*, encoding a conserved hypothetical protein with unknown function. The sequence of each gene was retrieved from 118 genome sequences, including those of 23 strains isolated from zucchini (Lacault et al. 2020), 14 strains isolated from cucurbits (Newberry et al. 2019), the two clade-2d strains isolated from zucchini seeds (this work) and 79 other PG2 strains (Supplementary Table 1), and aligned. Primers were designed in conserved regions in order to amplify and sequence polymorphic fragments. The primer pairs [Psyr3208-F ^5’^ATGATGCTGGTGCCGATGTA^3’^ and Psyr3208-R ^5’^ATCAGCGAGTAGACAGAACC^3’^], [Psyr3420-F ^5’^ATGGTGTTCATCGCCGTGACC^3’^ and Psyr3420-R ^5’^TCGCCTTCGACACCCAGCAG^3’^], and [Psyr4880-F ^5’^ATGCACGGCTCCGAACGAAA^3’^ and Psyr4880-R ^5’^AGTCTCCAGCGCCCGTCCAC^3’^] allowed the amplification of 883-pb, 343-pb, and 416-bp fragments, respectively. Supplementary primers, Psyr3208-MF ^5’^TCGGCCGAGTTCGCACCGAT^3’^ and Psyr3208-MR ^5’^CTGCCCGAGTAGATACCGAT^3’^ were designed in *Psyr3208* to improve sequencing. PCRs were performed in a 50-µL volume containing 10 µL of buffer, 0.4 U.µL^-1^ of Taq polymerase (GoTaq, Promega), 200 µM of dNTPs, 0.5 µM of each primer and 5 µL of boiled bacterial suspension. For *Psyr3208*, amplification conditions were 5 min at 94°C followed by 20 cycles of 30 s at 94°C, 30 s at 60°C (with a 0.5°C decrease at each cycle) and 1 min at 72°C, then 15 cycles of 30 s at 94°C, 30 s at 50°C and 1 min at 72°C. For *Psyr3420* or *Psyr4880*, PCR conditions were 5 min at 94°C followed by 35 cycles of 30 s at 94°C, 30 s at 62°C (for *Psyr3420*) or 66°C (for *Psyr4880*) and 1 min at 72°C. A final step of 10 min at 72°C was added to achieve elongation for all PCR tests. Amplified fragments were Sanger-sequenced with F- and R-primers, and, in addition with MF- and MR-primers for *Psyr3208* (Genoscreen, France). Geneious version 9.1.7 software was used to assemble, align, orientate, and trim sequences according to the reading frame. Sequences of partial CDSs were deposited in Genbank under the accession numbers MW892843-MW892968 and were 825-bp (*Psyr3208*), 318-bp (*Psyr3420*) et 309-bp (*Psyr4880*) long. Genes were amplified in strains whose genome sequence was not available. Homologous sequences retrieved from genome sequences (Table 2) were added to the data set.

**Table 2.** Genomic sequences used for MLSA.

| Strain code<sup>a</sup> | Country of isolation<sup>b</sup> | Host of isolation | Year of isolation<sup>b</sup> | Phylogroup, clade and cluster<sup>c</sup> | BIOSAMPLE |
| --- | --- | --- | --- | --- | --- |
| ICMP 2844<sup>PT</sup> (pv. *tomato*) | UK | *Lycopersicon esculentum* | 1960 | 1a | SAMN03976246 |
| Pmp 19322<sup>PT</sup> (pv. *morsprunorum*) | USA | *Prunus domestica* | na | 1b | SAMN03976519 |
| ICMP 3923<sup>PT</sup> (pv. *theae*) | Japon | *Thea sinensis* | 1970 | 1b | SAMN04033035 |
| Cit7 (*P. syringae*) | na | *Citrus sinensis* | na | 2a | SAMN02471314 |
| 58T<sup>T</sup> (*P. cerasi*) | Poland | *Prunus cerasus* | 2007 | 2a | SAMEA3894894 |
| ICMP 4048<sup>PT</sup> (pv. *papulans*) | Canada | *Malus pumila* | 1973 | 2a | SAMN03976275 |
| 03-19A | Florida | *Cucumis melo* | 2003 | 2a-D | SAMN06263767 |
| 200-1 | Georgia | *Cucurbita* sp. | 2000 | 2a-D | SAMN06318380 |
| P77 | France | *Cucurbita pepo* | 2010 | 2a-D | SAMN12288854 |
| P73 | USA | *C. pepo* | 2010 | 2a-D | SAMN12288853 |
| P78 | France | *C. pepo* | 2010 | 2a-D | SAMN12288860 |
| P87 | China | *C. pepo* | 2011 | 2a-D | SAMN12289083 |
| BS2121 | California | *Cucurbita* sp. | 2006 | 2a-E | SAMN06317958 |
| ZUM3984 | China | *Cucurbita* sp. | 2008 | 2a-E | SAMN06318038 |
| CFBP 8692 (P66) | Thailand | *C. pepo* | 2010 | 2a-E | SAMN12305548 |
| P89 | France | *C. pepo* | 2011 | 2a-E | SAMN12289141 |
| 13-139B | Florida | *Citrullus lanatus* | 2013 | 2a-F | SAMN06317916 |
| 13-429 | Florida | *C. lanatus* | 2013 | 2a-F | SAMN06318302 |
| ICMP 459<sup>PT</sup> (pv. *aptata*) | USA | *Beta vulgaris* | 1959 | 2b | SAMN03976295 |
| ICMP 3023<sup>T</sup> (pv. *syringae*) | UK | *Syringa vulgaris* | 1951 | 2b | SAMN03976255 |
| NCPPB 4273<sup>PT</sup> (pv. *coryli*) | Italia | *Corylus avellana* | 1995 | 2b | SAMN03481961 |
| P108 | Chile | *C. pepo* | 2011 | 2b | SAMN12288112 |
| 1188_1 | California | *C. pepo* | nd | 2b | SAMN03976479 |
| HS191 (pv. *syringae*) | Australia | *Panicum miliaceum* | 1969 | 2ba | SAMN03267749 |
| CFBP 8693 (P99) | France | *C. pepo* | 2005 | 2ba-A | SAMN12305539 |
| P12832 | France | *C. pepo* | 2006 | 2ba-A | SAMN12284164 |
| P12854 | France | *C. pepo* | 2002 | 2ba-A | SAMN12284309 |
| P12855 | Chile | *C. pepo* | 2007 | 2ba-A | SAMN12284728 |
| P113 | Thailand | *C. pepo* | 2011 | 2ba-A | SAMN12288386 |
| P118 | France | *C. pepo* | 2013 | 2ba-A | SAMN12288431 |
| P121 | Chile | *C. pepo* | 2013 | 2ba-A | SAMN12288432 |
| P123 | China | *C. pepo* | 2013 | 2ba-A | SAMN12288472 |
| P127 | India | *C. pepo* | 2013 | 2ba-A | SAMN12288592 |
| P139 | China | *C. pepo* | 2015 | 2ba-A | SAMN12288625 |
| P22 | USA | *C. pepo* | 2009 | 2ba-A | SAMN12288683 |
| P5 | France | *C. pepo* | 2008 | 2ba-A | SAMN12288805 |
| P79 | France | *C. pepo* | 2010 | 2ba-A | SAMN12289058 |
| P84 | China | *C. pepo* | 2010 | 2ba-A | SAMN12289072 |
| P12831 (co7A) | France | *C. pepo* | 2005 | 2ba-A | SAMN12289147 |
| PS711 | Serbia | *Cucurbita* sp. | 2013 | 2ba-A | SAMN10486866 |
| 13-509A | Florida | *Cucurbita* sp. | 2013 | 2ba-A | SAMN06263769 |
| KL004-k1 | Australia | *C. pepo* | 2016 | 2ba-A | SAMN23013829 |
| 13-C2 | Florida | *C. lanatus* | 2013 | 2ba-B | SAMN06263768 |
| 13-140A | Florida | *C. lanatus* | 2013 | 2ba-B | SAMN06256456 |
| 14-410 | Florida | *C. lanatus* | 2014 | 2ba-B | SAMN06270325 |
| 14-32 | Florida | *C. lanatus* | 2014 | 2ba-B | SAMN06257475 |
| 14-Gil | Florida | *C. lanatus* | 2014 | 2ba-B | SAMN06317944 |
| CC457 (*P. syringae*) | France | *C. melo* | na | 2ba-B | SAMN02471568 |
| P12857 | France | *C. pepo* | 2007 | 2ba-B | SAMN12288124 |
| ZUM3584 | Italy | *Cucurbita* sp. | 2005 | 2ba-C | SAMN06318381 |
| KFR003-1 | Australia | *C. pepo* | 2016 | 2ba-C | SAMN23013828 |
| 77-4C | Australia | *C. pepo* | 2016 | 2ba-C | SAMN23013827 |
| ICMP 19117<sup>PT</sup> (*P. congelans*) | Germany | *Poaceae* | 1994 | 2c | SAMN03976302 |
| B728A (pv. *syringae*) | USA | *Phaseolus vulgaris* | na | 2d | SAMN09235866 |
| ICMP 16925<sup>PT</sup> (pv. *solidagae*) | Japon | *Solidago altissima* | na | 2d | SAMN03976256 |
| ICMP 2802<sup>PT</sup> (pv. *aceris*) | USA | *Acer* sp. | na | 2d | SAMN03976301 |
| 1212 (pv. *syringae*) | UK | *Pisum sativum* | na | 2d | SAMN02471561 |
| HRI-W 7872 (pv. *syringae*) | UK | *Prunus domestica* | 2000 | 2d | SAMN03992220 |
| P129 | China | *C. pepo* | 2013 | 2d | SAMN18647612 |
| P135 | India | *C. pepo* | 2014 | 2d | SAMN18647613 |
| ICMP 2740<sup>PT</sup> (pv. *phaseolicola*) | Canada | *P. vulgaris* | 1949 | 3 | SAMN03976265 |
| NM002 (pv. *lachrymans*) | China | *Cucumis sativus* | 1984 | 3 | SAMN06606132 |
| LMG 5060<sup>PT</sup> (*P. coronafaciens*) | UK | *Avena sativa* | 1958 | 4 | SAMN03144492 |
| ICMP 2823<sup>PT</sup> (pv. *cannabina*) | Hongria | *Cannabis sativus* | 1957 | 5 | SAMN05216597 |
| ICMP 4091<sup>PT</sup> (pv. *tagetis*) | Zimbabwe | *Tagetes erecta* | 1972 | 6 | SAMN03976254 |
| ICMP 3272 (*P. viridiflava*) | New Zeland | *Actinidia deliciosa* | 1971 | 8 | SAMN03976434 |

<sup>a</sup> T = type strain; <sup>PT</sup> = pathotype strain <sup>b</sup> na = not available <sup>c</sup> phylogroup, clade and cluster according to Berge et al 2014, Newberry et al 2019 and Lacault et al 2020.

### MLSA of the strain collection

The sequences of *gapA, gltA, gyrB,* and *rpoD* (Lacault et al. 2020), encoding the glyceraldehyde-3-phosphate dehydrogenase A, the citrate synthase A, the B subunit of DNA gyrase, and the RNA polymerase sigma70 factor RpoD, respectively, were concatenated to the sequences of *Psyr3208*, *Psyr3420*, and *Psyr4880,* to obtain a 3,459 bp fragment. A maximum likelihood tree was generated with MEGA 7 (Kumar et al. 2016). The best fit model was the Tamura-Nei model using the discrete Gamma distribution with invariant sites (G + I) and 1000 bootstrap replicates were used.

### Genome sequencing and analyses

The genomes of strains P129_2d_ and P135_2d_ belonging to the clade 2d were sequenced. Their DNA was extracted using the Wizard Genomic DNA Purification Kit (Promega) and was paired-end sequenced (2 × 150 bp) using the Illumina HiSeq X-Ten at Beijing Genomics Institute (China). Genome assembly was performed with SOAPdenovo version 2.04, SOAPGapCloser version 1.12 (Luo et al. 2012), and Velvet version 1.2.02. (Zerbino and Birney 2008). Fasta-formatted assemblies were deposited in NCBI with accession numbers SAMN18647612 and SAMN186476113. Phylogenic position of clade-2d strains was determined by the percentage of shared 15-mers (Briand et al. 2019) on the dataset of 118 genomes (Supplementary Table 1). Percentage of shared k-mers and distance matrix were calculated with Ki-S as were dendrograms (https://iris.angers.inra.fr/galaxypub-cfbp) (Briand et al. 2019). T3E repertoire of each strain was predicted through tBLASTN search (identity higher than 80% on at least 80% of CDS length) on 138 amino acid sequences of T3Es (Hopsdatabase at http://www.pseudomonas-syringae.org/). Putative frameshift mutations were checked by aligning each T3E sequence with the corresponding curated reference sequence with MultAlin (http://multalin.toulouse.inra.fr/multalin/) (Corpet 1988).

### Spiking of seed samples and analysis of seed lots

Healthy seed samples (negative result for a 2,000-seed analysis) of *C. pepo* subsp. *pepo* cv. Lorea (500 seeds) were soaked in 3 mL.g^-1^ of seeds of phosphate-buffered saline (PBS, Sigma) plus Tween 20 (0.02% v/v) and shaken (105 rpm) during two hours at room temperature. A fresh bacterial suspension calibrated at 1 × 10^8^ CFU.mL^−1^ was diluted with seed extract solutions in order to end up with spiked suspensions at concentrations ranging from 1 to 1 × 10^7^ CFU.mL^-1^, which were confirmed by dilution plating. One mL per dilution was centrifugated 5 min at 13,000 g. Supernatant was discarded and DNA was extracted using the QuicPick^TM^ SML Plant DNA purification kit according to the supplier’s instructions (Bio-Nobile, Turku, Finland) in an automated system (Caliper Zephyr, PerkinElmer). DNA was eluted from magnetic beads using 30 µL of elution buffer. DNAs of bacterial suspensions in sterile distilled water were also extracted and used as standards or controls. Dilution series of DNA solutions ranged from 1 to 1 × 10^7^ CFU.mL^-1^. One to five µL of DNA were used per qPCR sample. Each sample was tested in triplicate. Four commercial zucchini seed lots (I to IV) were tested in samples of 500 seeds or in five subsamples of 100 or 10 seeds. One µL of DNA was used per qPCR sample. Subsampling allowed the determination of the contamination rate of the seed lot according to Maury et al. (1985) when subsamples had the same size or according to Swaroop (1951) for degressive subsamplings.

### Identification of targets and primer design

Specific sequences of clades and clusters were searched for in the genomic sequences using the SkIf_with_DSK tool (Briand et al. 2016; https://iris.angers.inra.fr/galaxypub-cfbp). SkIf_with_DSK identifies specific k-mers within a group of genome sequences (in-group) that are absent in the other genome sequences (out-group). It also provides their precise locations on a reference genome and uses the positions to concatenate the overlapping k-mers into long-mers (Denancé et al. 2019). A k-mer size of 22 was used to search for specific sequences using the dataset of 118 genome sequences (Supplementary Table 1). Specific long-mers were searched for in clusters 2ba-A, 2ba-B, 2ba-C, 2a-D and 2a-E, in larger groups 2ba-ABC and 2a-DE encompassing the clusters of VCZ strains, and in clades 2b and 2d (Figure 2 and Supplementary Table 1). Different tree nodes were tested to isolate long-mers specific of clade-2b strains including or not pathovar *pisi* or the clade 2ba (Figure 2). Precise composition of in- and out-groups per marker is given in Supplementary Table 1. In order to design PCR tests and qPCR tests, only long-mers with a size greater than or equal to 70 bp were considered.

Primer3 2.3.4 was used to select primers in specific long-mers for all classical PCRs with parameters set up with an optimal primer size of 20 bp, a product length from 150 up to 350 bp, and a Tm of 60°C (± 3°C). For qPCR tests, target sequences (*avrRpt2*, *hopZ5* and long-mers specific of clade 2d) were extracted from genome sequences using the bioinformatic tool Extract_Genes_Genomes (https://iris.angers.inra.fr/galaxypub-cfbp) and aligned using Geneious R9. Primers and probe combinations for qPCR assays were designed within these specific regions using Geneious 9.1.8 (Koressaar and Remm, 2007). The parameters were set up with an optimal primer size of 20 bp, an optimal product size of 80 bp, and Tm of 60°C and 70°C for primers and probe, respectively. Primer3, release 2.4.0 was used to design qPCR for clade-2b strains. Dimer- and hairpin-forming primers and probes were discarded.

### *In silico* evaluation of the specificity of the PCR and qPCR tests

Specificity of primers and associated probes was tested *in silico* using PrimerSearch (Val Curwen, Human Genome Mapping Project, Cambridge, UK), first, on our collection of 121 genomic sequences, and second, on the 194,438 whole Genome Shotgun (WGS) sequences of Bacteria and Archae available in the NCBI database (03/2019 release). When necessary, phylogenetic position of strains was checked using KISonWGS (Briand et al. 2020) (https://iris.angers.inra.fr/galaxypub-cfbp). KISonWGS identifies WGS neighbor genome sequences of query input genome file. For multiplex qPCRs, sets of primers and probe and combinations of primers and probes were tested on the genomes of P99_2ba-A_, P12857_2ba-B_, ZUM3584_2ba-C_, P77_2a-D_, P66_2a-E_, 13-139B_2a-F_, P108_2b_ and P135_2d_ using AmplifX version 2.0.7 (Nicolas Jullien, Aix-Marseille Univ, CNRS, INP, Inst Neurophysiopathol, Marseille, France; https://inp.univ-amu.fr/en/amplifx-manage-test-and-design-your-primers-for-pcr) to check absence of dimers and cross-amplifications.

### PCR and qPCR assays, specificity and sensitivity

Classical PCRs were performed in a 20-µL volume containing 200 nM of each primer, 200 nM of dNTP, 0.08 µL of GoTaq2 and 4 µL of its 5x corresponding buffer (Promega), and 5 µL of boiled calibrated bacterial suspensions. Amplification reactions were run in a thermocycler 5 min at 95°C, 35 cycles of 30 s at 94°C, 30 s at 57°C, and 1 min at 72°C, followed by 3 min at 72°C. Five µL of sample were used for electrophoresis 20 min at 100 V in 1% agarose gel stained with ethidium bromide, and visualized on a UV transilluminator. All qPCRs were performed in a Bio-Rad CFX96 Touch thermocycler and analysed with Bio-Rad CFX Manager 3.1 software. Each individual and multiplexed TaqMan reaction was performed in a final volume of 10 µL containing 5 µL of Sso Advanced Universal Probes Supermix (Bio-Rad), 600 nM of each primer, 200 nM of probe, and 1 µL of boiled calibrated bacterial suspension. The amplification program was 3 min at 95°C for and 40 cycles of 15 s at 95°C and 30 s at 60°C. The qPCR on *sylC* was also performed in a final volume of 20 µL containing 10 µL of MasterMix buffer (Eurogentec, Belgium), 900 nM of each primer, 250 nM of probe, with 5 µL of boiled calibrated bacterial suspensions. The amplification program was 10 min at 95°C for and 40 cycles of 15 s at 95°C and 15 s at 60°C. For each reaction, equation curve, efficiency of qPCR (E = 10^(−1/slope)^) and the correlation coefficient R^2^ were determined using calibrated DNA solutions (1 µg.mL^-1^ to 1 pg.mL^-1^) of strains P99 for qPCR on *sylC* and *avrRpt2* (primers and probes 4Ba and Rpt2, respectively), P66 for qPCR on *hopZ5* (primers and probe Z5) and P108 and P129 for qPCR for clades 2b* and 2d (primers and probes 2b* and 2d, respectively).

The specificity of the primers and probes was tested *in vitro* with an appropriate collection of target and non-target bacterial strains (Table 1). Specificity was checked in simplex and multiplex at least twice in duplicates using boiled bacterial suspensions (OD_650 nm_ = 0.1). Sensitivity of the multiplex qPCR assays was evaluated at least twice in triplicates for boiled bacterial suspensions (10^8^ to 10^3^ CFU.mL^-1^) and DNAs extracted from 1 mL of seed extracts spiked with bacterial suspensions for final concentrations ranging from 10^7^ to 10^1^ CFU.mL^-1^. The limit of detection (LOD) was the lowest concentration with a positive signal in, at least, two out of the triplicates.

## Results

### Characterization of the strain collection using the seven-genes MLSA scheme

The analysis of our strain collection using the 7-genes MLSA scheme revealed that 29 strains belonged to cluster 2ba-A, seven to cluster 2ba-B, two to cluster 2ba-C, nine to cluster 2a-D and six to cluster 2a-E (Figure 1). VCZ strains were distributed in five clusters that all included strains responsible for other diseases on cucurbits (Figure 1). No VCZ strain belonged to the cluster F. The strains of clade 2b and 2d that were analyzed in MLSA, were genetically diverse and did not form any clusters.

**Figure 1.**
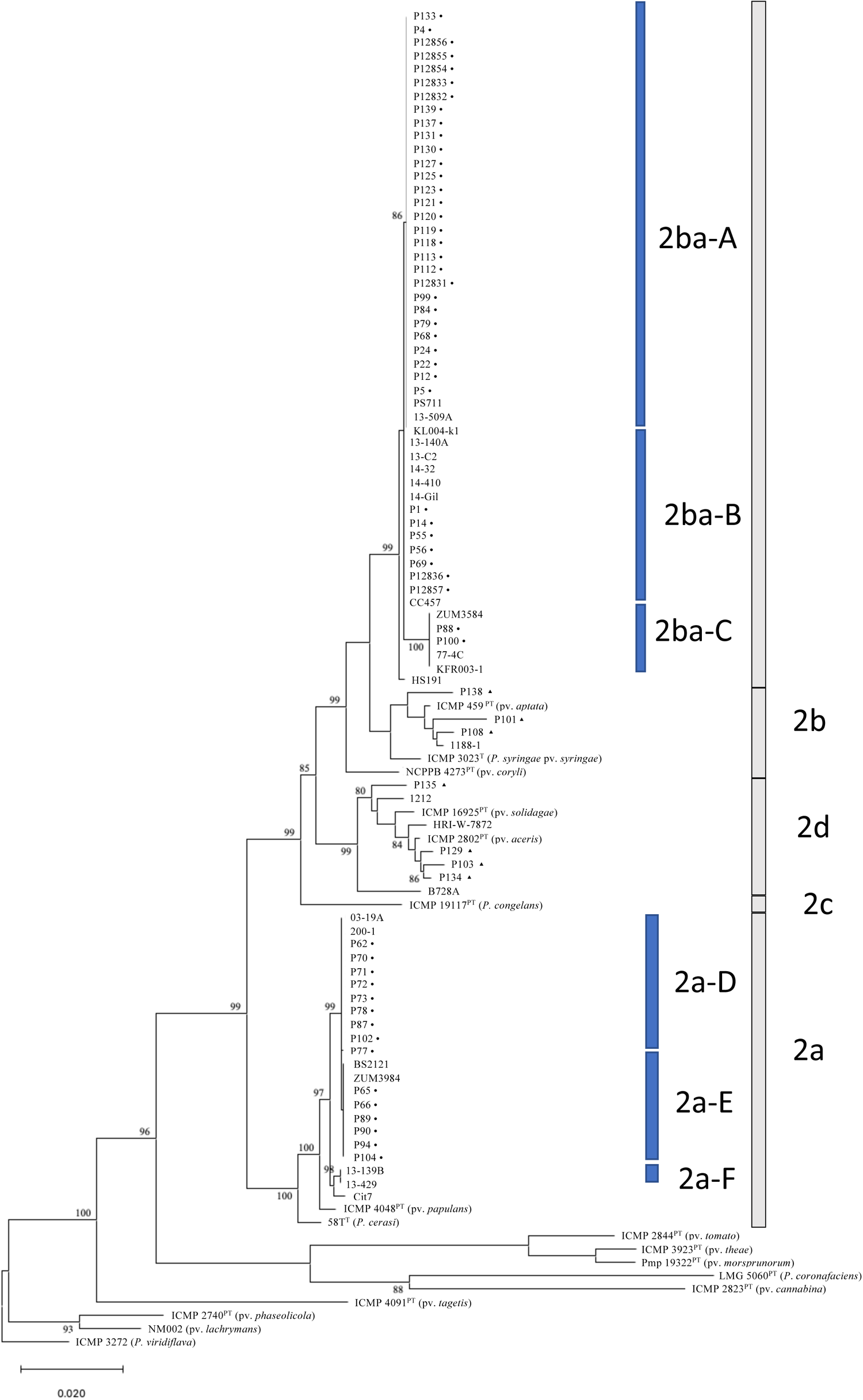
Phylogenetic relationships of strains isolated from seed lots between 2002 and 2014. Maximum likelihood (ML) tree was based on concatenated partial sequences of *gapA, gltA, gyrB*, *rpoD*, *Psyr3208, Psyr3420* and *Psyr4880* (3,405 bp). Bootstrap scores (1,000 replicates) are displayed at each node. Clades are indicated based on several studies (Berge et al. 2014; Bull and Koike 2015; Newberry et al. 2019) and clusters according to Lacault and colleagues (2020). Black symbols indicate strains from our collection: circles (VCZ strains) and triangles (clade-2b and −2d strains isolated from zucchini seeds).

**Figure 2.**
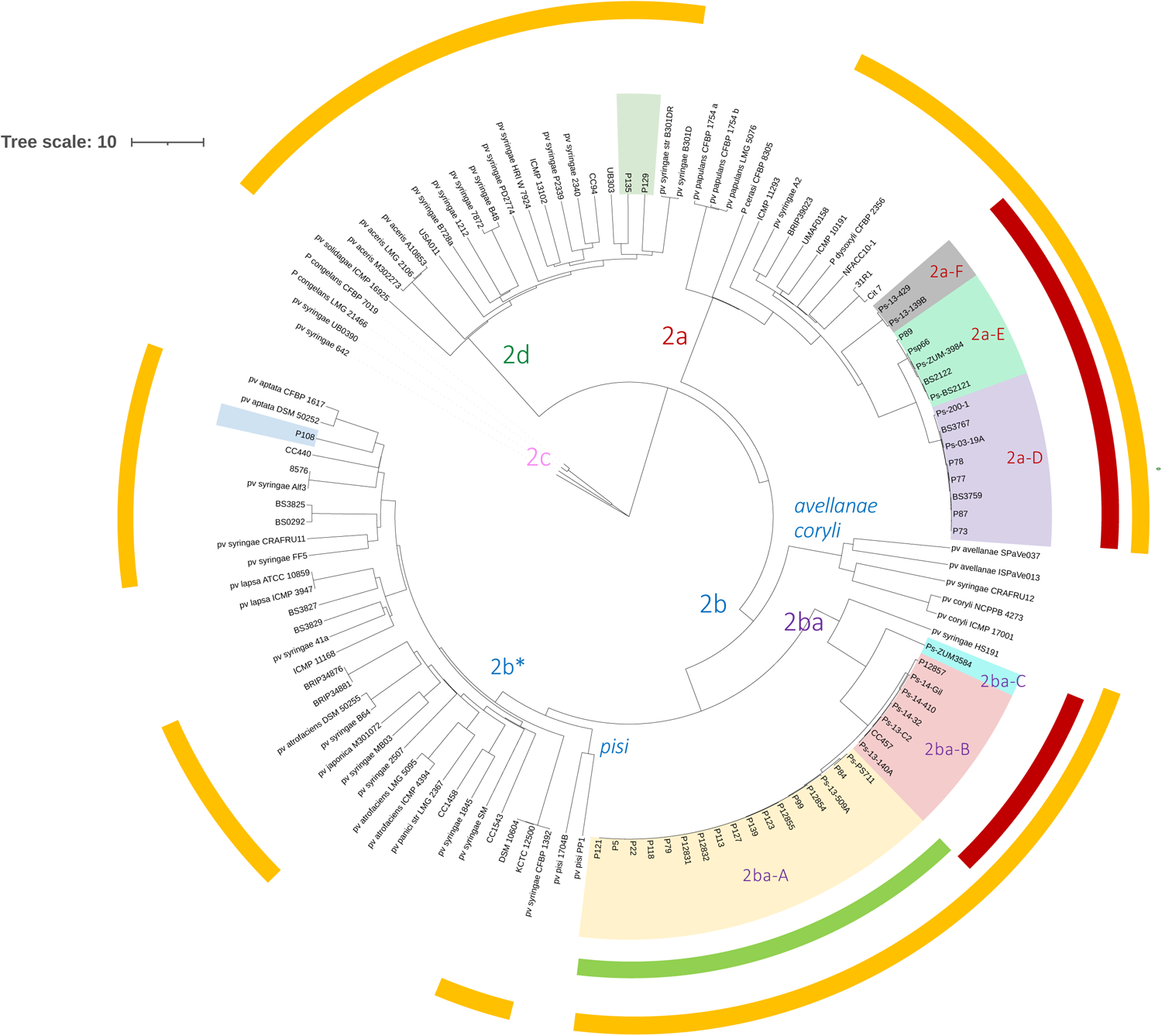
Phylogenetic groups chosen for PCR design. Dendrogram built on distances calculated from shared k-mer matrices obtained with KIS on a dataset of 118 genomic sequences of *Pseudomonas syringae* phylogroup 2. Clades are indicated based on several studies (Berge et al. 2014; Bull and Koike 2015; Newberry et al. 2019) and clusters according to Lacault and colleagues (2020). Presence of *avrRpt2, hopZ5,* and *sylC* is indicated by a green, red and yellow color on the outside circles, respectively.

### Insights on clade-2d strains

Genomes of strains P129_2d_ and P135_2d_ isolated from zucchini seed lots were sequenced to be used as reference for clade 2d and the percentage of shared k-mers between their sequences confirmed this phylogenetic position (Figure 2), as well as their identification based on MLSA (Figure 1). Genome sequences of strains P129_2d_ and P135_2d_ contained 11 and 12 T3E encoding genes, respectively, among which *avrE1, hopAA1, hopAE1, hopAG1, hopAH1, hopBC1, hopH1, hopI1,* and *hopM1* that were also present in most VCZ strains (Lacault et al. 2020). The T3E genes *hopA2* and *hopAY1* were present in both strains whereas *hopBE1* was only present in P135_2d_. These clade-2d strains lacked *hopAZ1, hopBA1,* and *hopA1* which are present in VCZ strains of clades 2a and 2ba (Lacault et al. 2020). The T3E genes *avrRpt2* and *hopZ5*, one of which is present in the genomic sequences of all VCZ strains, were not present in P129_2d_ and P135_2d_ genome sequences. Similarly, the clade-2b singleton P108_2b_ has a small T3E repertoire and lacks *hopA1, avrRpt2* and *hopZ5* (Lacault et al. 2020).

### Design of primers and probes on identified targets

Specific sequences of each of the clusters 2ba-A to 2a-E, of larger groups 2ba-ABC (including clusters 2ba-A, -B, and -C), 2a-DE (including clusters 2a-D and -E), and of clades 2b and 2d (Figure 2), were identified in genome sequences using SkIf_with_DSK. Among the 82 long-mers specific of cluster 2ba-A, the 45 of cluster 2ba-B, the 147 of cluster 2ba-C, the 124 of cluster 2a-D, the 213 of cluster 2a-E, the 239 of the group 2ba-ABC, the 377 of the group 2a-DE, and the 2554 of the clade 2d, the first sequence of each targeted group was generally used as a template sequence for PCR primers or qPCR primers and probe designs (Table 3). Concerning clade 2b, only two specific long-mers allowing a qPCR primers and probe design, were identified for the node 2b*, excluding branches encompassing clade 2ba and pathovars *coryli*, *avellanae* and *pisi* (Figure 2, Table 3). No specific qPCR primers could be designed for the other clade-2b nodes, because of (i) the lack of specific long-mers, (ii) long-mers were not appropriate to design a test, or (iii) the test was not inclusive of clade-2b strains isolated from zucchini (data not shown). Primers and probes for Taqman qPCR were designed within *avrRpt2* and *hopZ5* (Table 3).

**Table 3.**
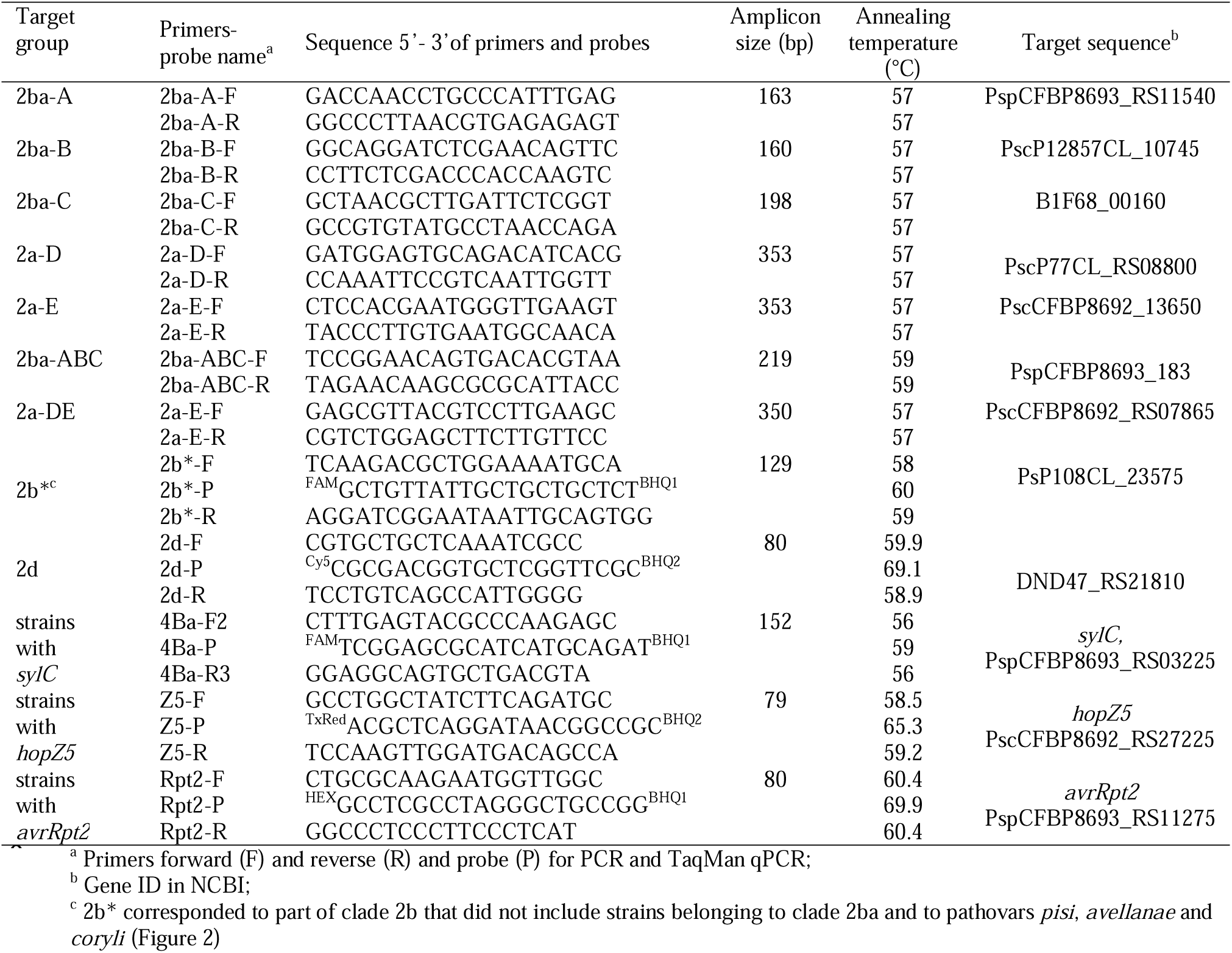
Primers and probes designed to detect and identify strains belonging to the species complex *P. syringae,* isolated from zucchini seeds

### *In silico* and *in vitro* evaluation of the specificity of the PCR and qPCR assays

The PCR tools developed to determine the position of strains in the different clusters and clades were all 100% inclusive *in vitro*. Inclusivity was also excellent *in silico*, as only one exception was identified for one strain in clade-2b* (Table 4). A limited number of cross-reactions was highlighted, giving an exclusivity of 99.96 to 100% depending on the test (Supplementary Table 2, and Tables 1 and 4). Several of these cross reactions involved strains with in-between genotypes, revealing a greater diversity than expected within 2ba clade. For the test targeting 2ba-A cluster, strain 1188-1 (SAMN03976479) isolated from zucchini, classified in clade 2b on the basis of its whole genome sequence, was positive *in silico* with the 2ba-A test (Table 4). This genome was positive for other PCR tests targeting cluster 2ba-A such as 2ba-ABC, 4Ba, and Rpt2 (targeting *avrRpt2*) PCR tests and matched also primers and probe for clade 2b (Supplementary Table 2). Similarly, positive reactions with 2ba-A and Rpt2 primers were observed *in silico* or *in vitro* for three strains belonging to clusters 2ba-B and 2ba-C2 (Supplementary Table 2 and Table 4). For the PCR test dedicated to cluster 2ba-B strains, three cross-reactions were observed *in silico* for *P. coronafaciens* genomes (PG4, Table 4), that were confirmed *in vitro* by a positive signal for the type strain of *P. coronafaciens* (Table 1). For cluster 2ba-C, primers annealed with 4 mismatches *in silico* with a strain of clade 2a and gave a band at the wrong size *in vitro* for one clade-2b strain. Two genomes of *P. coronafaciens* perfectly matched with 2a-D primers. Primers for group 2ba-ABC annealed *in silico* with mismatches on two strains, of which strain 1188-1. Primers for group 2a-DE annealed the two cluster 2a-F genome sequences with three mismatches. Concerning 2b* PCR test, the clade-2b genome sequence of strain ICMP 4917 (SAMN03976350) did not match *in silico*. Numerous *in silico* unperfect matches were obtained for primers and probe targeting clade 2b* on non-targeted genomes but they were not confirmed *in vitro* on representative strains. Genome sequence of strain ICMP 4996 (SAMN03976403), named *P. coronafaciens* pv. *striafaciens* in the NCBI database, was classified in the 2b clade according to our KISonWGS assignation and to Dillon and colleagues (Dillon et al. 2019b) and not in PG4 as expected by its taxonomic assignation. This genomic sequence perfectly matched the 2b* primers and probe. This result was confirmed *in vitro* with the synonymous strain CFBP 1750 that gave a positive signal with this test. This particular case was considered a problem of misclassification and not a real cross reaction. Primers and probe dedicated to clade-2d strains gave one cross reaction *in silico* with the clade-2b strain 3023 (SAMN05861187).

**Table 4.**
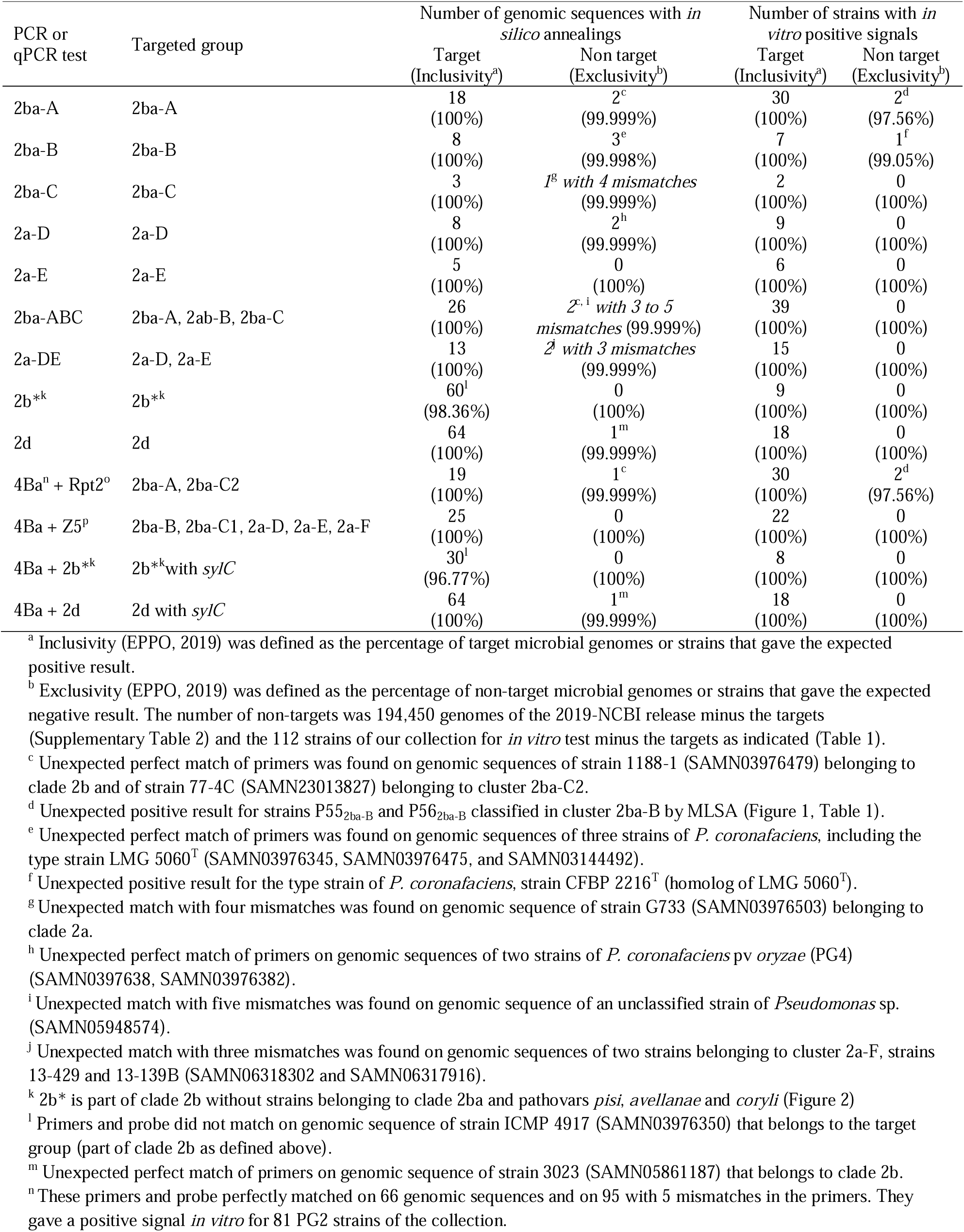
*In silico* and *in vitro* specificity of primers and probes.

*In silico* and *in vitro* testing 4Ba qPCR test confirmed that this test was 100% inclusive of VCZ strains but detected also some other PG2 strains despite mismatches in the primers (Table 1 and Supplementary Table 2). Primers and probe targeting the gene *avrRpt2* annealed on the genomic sequences of all clusters 2ba-A and 2ba-C2 strains, strain 1188-1_2b_ and some strains of pathovars *tomato, spinacea, maculicola* and *tagetis*. These *in silico* results were confirmed *in vitro* with the detection of one strain of pathovar *tomato* and another one of pathovar *spinacea*. The Z5 primers and probes (targeting *hopZ5*) detected *in silico* and *in vitro* strains of clusters 2ba-B and -C and 2a-D, -E and -F, and strains of the pathovar *actinidiae*. This later pathovar is known to have *hopZ5*. Unperfect matches were found *in silico* for pathovars *berberidis*, *delphinii,* and *phaseolicola*, and *in vitro* tests of representative strains were negative. All strains of those cross reacting pathovars were negative with the 4Ba qPCR test.

### Development of a multiplex qPCR test to identify VCZ strains

A multiplex TaqMan qPCR test combining primers and probes targeting *sylC*, the two genes encoding the pathogenicity factors AvrRpt2 and HopZ5 and the clade-2d strains was developed to detect all VCZ strains and gain information on contaminating strains, in particular on their pathogenicity. All targeted sequences, when present in a genome, were found in single copy. Individually the three PCR tests targeting genes encoding the pathogenicity factors SylC, AvrRpt2 and HopZ5 were not specific for VCZ strains (Supplementary Table 2). However, VCZ strains from main clusters are characterized by the simultaneous presence of *sylC* and either *avrRpt2* or *hopZ5* (Figure 2 and Table 1), as *avrRpt2* characterizes 2ba strains that have a narrow host range on cucurbits whereas *hopZ5* is present in strains with a wider host range (Lacault et al. 2020; Djitro et al. 2022a). Clade-2b* and −2d strains were also isolated from zucchini seeds and gave a positive signal with 4Ba test (targeting *sylC*) justifying the need to include their corresponding qPCR tests in the multiplex. However, the attempt to add the clade 2b* qPCR test in the multiplex was abandoned due to qPCR quantitative parameters incompatible with other tests (data not shown).

Specificity of each qPCR test was not affected by multiplexing. *In silico* simulation of multiplex qPCR test with AmplifX did not show any cross reactions between the different pairs of primers and the probes. The efficiencies of the qPCR tests were evaluated in simplex and multiplex on serial dilutions of DNA extracted from each target strain. Parameters such as PCR efficiency and R^2^ of regression curves were correct in both simplex and multiplex, except for 4Ba test whose efficiency was better in the conditions of multiplex than simplex qPCR (Table 5). Specificity of each qPCR test was identical in multiplex and in simplex (data not shown).

**Table 5.** Efficiencies of simplex and multiplex qPCR tests on calibrated DNA extracts

| qPCR test<br>(strain) | DNA<br>concentration | Number of genome<br>copies.mL<sup>-1a</sup> | Mean Cq value (SEM)<sup>b</sup> |  |
| --- | --- | --- | --- | --- |
|  |  |  | Simplex | Tetraplex |
| 4Ba<br>(P99<sub>2ba-A</sub>) | 1 µg.mL<sup>-1</sup> | 1.65 × 10<sup>8</sup> | 22.43 (0.29) | 19.43 (0.09) |
|  | 100 ng.mL<sup>-1</sup> | 1.65 × 10<sup>7</sup> | 26.05 (0.13) | 22.85 (0.09) |
|  | 10 ng.mL<sup>-1</sup> | 1.65 × 10<sup>6</sup> | 29.13 (0.07) | 26.36 (0.12) |
|  | 1 ng.mL<sup>-1</sup> | 1.65 × 10<sup>5</sup> | 32.69 (0.39) | 29.37 (0.05) |
|  | 100 pg.mL<sup>-1</sup> | 1.65 × 10<sup>4</sup> | 37.98 (1.93) | 33.26 (0.46) |
|  | 10 pg.mL<sup>-1</sup> | 1.65 × 10<sup>3</sup> | 38.95 (0.26) | NA<sup>e</sup> |
|  | 1 pg.mL<sup>-1</sup> | 1.65 × 10<sup>2</sup> | NA | NA |
| Efficiency |  |  | 82 | 94.9 |
| R2 |  |  | 0.963 | 0.989 |
| slope |  |  | -3.84 | -3.451 |
| Rpt2<br>(P99<sub>2ba-A</sub>) | 1 µg.mL<sup>-1</sup> | 1.65 × 10<sup>8</sup> | 21.55 (0.15) | 19.77 (0.08) |
|  | 100 ng.mL<sup>-1</sup> | 1.65 × 10<sup>7</sup> | 25.11 (0.12) | 23.19 (0.08) |
|  | 10 ng.mL<sup>-1</sup> | 1.65 × 10<sup>6</sup> | 28.49 (0.12) | 26.64 (0.10) |
|  | 1 ng.mL<sup>-1</sup> | 1.65 × 10<sup>5</sup> | 32.47 (0.17) | 30.01 (0.16) |
|  | 100 pg.mL<sup>-1</sup> | 1.65 × 10<sup>4</sup> | 35.47 (0.21) | 33.93 (0.35) |
|  | 10 pg.mL<sup>-1</sup> | 1.65 × 10<sup>3</sup> | 39.05 (0.31) | NA |
|  | 1 pg.mL<sup>-1</sup> | 1.65 × 10<sup>2</sup> | 21.55 (0.15) | NA |
| Efficiency |  |  | 92.9 | 92.3 |
| R2 |  |  | 0.997 | 0.99 |
| slope |  |  | -3.505 | -3.521 |
| Z5<br>(P66<sub>2a-E</sub>) | 1 µg.mL<sup>-1</sup> | 1.62 × 10<sup>8</sup> | 21.18 (0.08) | 18.34 (0.12) |
|  | 100 ng.mL<sup>-1</sup> | 1.62 × 10<sup>7</sup> | 24.20 (0.05) | 22.25 (0.27) |
|  | 10 ng.mL<sup>-1</sup> | 1.62 × 10<sup>6</sup> | 27.54 (0.12) | 25.36 (0.12) |
|  | 1 ng.mL<sup>-1</sup> | 1.62 × 10<sup>5</sup> | 30.95 (0.04) | 28.79 (0.17) |
|  | 100 pg.mL<sup>-1</sup> | 1.62 × 10<sup>4</sup> | 34.22 (0.24) | 33.11 (0.01) |
|  | 10 pg.mL<sup>-1</sup> | 1.62 × 10<sup>3</sup> | 36.68 (0.67) | 35.91 |
|  | 1 pg.mL<sup>-1</sup> | 1.62 × 10<sup>2</sup> | NA | NA |
| Efficiency |  |  | 95.8 | 90.2 |
| R2 |  |  | 0.991 | 0.995 |
| slope |  |  | -3.426 | -3.582 |
| 2d<br>(P129<sub>2d</sub>) | 1 µg.mL<sup>-1</sup> | 1.61 × 10<sup>8</sup> | 26.27 (0.09) | 19.99 (0.07) |
|  | 100 ng.mL<sup>-1</sup> | 1.61 × 10<sup>7</sup> | 28.15 (0.41) | 24.25 (0.16) |
|  | 10 ng.mL<sup>-1</sup> | 1.61 × 10<sup>6</sup> | 32.03 (0.28) | 27.66 (0.10) |
|  | 1 ng.mL<sup>-1</sup> | 1.61 × 10<sup>5</sup> | 35.91 (0.61) | 30.49 (0.27) |
|  | 100 pg.mL<sup>-1</sup> | 1.61 × 10<sup>4</sup> | 38.08 (1.01) | 33.36 (0.29) |
|  | 10 pg.mL<sup>-1</sup> | 1.61 × 10<sup>3</sup> | 37.83 | 37.09 |
|  | 1 pg.mL<sup>-1</sup> | 1.61 × 10<sup>2</sup> | NA | NA |
| Efficiency |  |  | 99.8 | 101 |
| R2 |  |  | 0.994 | 0.988 |
| slope |  |  | -3.326 | -3.299 |

<sup>a</sup> The theoretical number of genome copies in DNA solutions was calculated with 1 pg corresponding to 978 Mb and genome sizes to 5.908, 6.185, and 6.044 Mb for P99<sub>2ba-A</sub>, P66<sub>2a-E</sub> and P129<sub>2d</sub>, respectively;
<sup>b</sup> NA = Not Amplified; values in *italics* correspond to a unique Cq value; mean Cq values were calculated on at least two positive values.

The LOD was defined as the lowest concentration giving a positive result in at least two out of the triplicates. The LOD of each qPCR test on calibrated DNA solutions in multiplex was 100 pg.mL^-1^ (0.1 pg per reaction corresponding to 16 genome copies), but was 10 times better in simplex for 4Ba, Rpt2 and Z5 tests (Table 5). Similarly, LODs in multiplex were 2.04 × 10^4^, 2.46 × 10^4^, 4 × 10^4^, and 5.80 × 10^3^ CFU.mL^-1^ (corresponding to 20, 25, 40 and 6 CFUs per reaction) were obtained with boiled calibrated bacterial suspensions for 4Ba, Rpt2, Z5 and 2d, qPCR respectively (Supplementary Table 3). Similar results were obtained in the second experiment, the apparent better LOD for 2d qPCR test with bacterial suspensions was observed in two independent experiments (Supplementary Table 3).

### Analysis of spiked and commercial seed lots with the newly developed multiplex qPCR test

Seed extracts were spiked with calibrated bacterial suspensions to determine the LOD of the multiplex qPCR test. Strains P99_2ba-A_, P66_2a-E_ and P135_2d_ were inoculated individually in different seed extracts (batches of 500 seeds soaked in buffer). For each strain, serial dilutions of a calibrated suspension were inoculated in different seed extract subsamples. Experiment was made independently twice with different inocula and seed extracts. A preliminary step of DNA extraction was performed to avoid potential PCR inhibitors present in seed extracts. DNAs were extracted from 1 mL samples and eluted in a 30-µL volume. By this process samples could potentially be concentrated 33-fold. Samples (1 µL per reaction) were amplified in triplicates with the multiplex qPCR test. The LODs of the spiked seed extracts corresponded to 2.46 × 10^3^, 2.46 × 10^3^, 3.22 × 10^3^, and 5.80 × 10^2^ CFU.mL^-1^ for 4Ba, Rpt2, Z5 and 2d, respectively (Table 6). The parameters of the standard curve on DNAs extracted from spiked seed-extracts (Table 6) were not different from those obtained on pure bacterial suspensions (Supplementary Table 4). These LODs were globally 10 times lower than the LODs obtained for calibrated bacterial suspensions (Table 5, Table 6, and Supplementary Table 4), due to the increase of DNA concentration during extraction and to the absence of inhibitors after DNA extraction. Thus, the protocol used to extract bacterial DNA seemed to be adapted to discard inhibitors from seed and recover bacterial DNA with a quality compatible with qPCR.

**Table 6.** Sensitivity of the multiplex qPCR test on seed extracts spiked with calibrated bacterial suspensions

| Strain | concentration<sup>a</sup><br>CFU.mL<sup>-1</sup> | First experiment |  | concentration<sup>a</sup><br>CFU.mL<sup>-1</sup> | Second experiment |  |
| --- | --- | --- | --- | --- | --- | --- |
|  |  | Mean Cq (SEM) |  |  | Mean Cq (SEM) |  |
|  |  | 4Ba | Rpt2 |  | 4Ba | Rpt2 |
| P99<sub>2ba-A</sub> | 2.46 × 10<sup>7</sup> | 18.16 (0.10) | 18.11 (0.06) | 3.28 × 10<sup>7</sup> | 18.18 (0.15) | 17.91 (0.23) |
|  | 2.46 × 10<sup>6</sup> | 21.94 (0.21) | 21.98 (0.17) | 3.28 × 10<sup>6</sup> | 21.48 (0.24) | 21.18 (0.18) |
|  | 2.46 × 10<sup>5</sup> | 25.53 (0.12) | 25.74 (0.09) | 3.28 × 10<sup>5</sup> | 25.19 (0.55) | 25.02 (0.31) |
|  | 2.46 × 10<sup>4</sup> | 28.01 (0.01) | 28.02 (0.05) | 3.28 × 10<sup>4</sup> | 26.67 (0.17) | 27.12 (0.04) |
|  | 2.46 × 10<sup>3</sup> | 31.75 (0.14) | 31.59 (0.36) | 3.28 × 10<sup>3</sup> | 32.5 (0.34) | 31.36 (0.50) |
|  | 2.46 × 10<sup>2</sup> | NA<sup>b</sup> | NA | 3.28 × 10<sup>2</sup> | NA | NA |
|  | 2.46 × 10<sup>1</sup> | NA | NA | 3.28 × 10<sup>1</sup> | NA | NA |
|  | 0 to 3 | NA | NA | 0 to 4 | NA | NA |
| Efficiency<sup>c</sup> |  | 99.9% | 100.9% |  | 100.7% | 102.9% |
| R<sup>2</sup> |  | 0.994 | 0.99 |  | 0.96 | 0.992 |
| Slope |  | -3.325 | -3.301 |  | -3.306 | -3.255 |
| P66<sub>2a-E</sub> | 3.22 × 10<sup>7</sup> | 20.68 (0.03) | 17.48 (0.06) | 4 × 10<sup>7</sup> | 21.76 (0.12) | 18.26 (0.06) |
|  | 3.22 × 10<sup>6</sup> | 24.81 (0.04) | 21.37 (0.03) | 4 × 10<sup>6</sup> | 25.61 (0.13) | 22.11 (0.08) |
|  | 3.22 × 10<sup>5</sup> | 27.51 (0.12) | 24.15 (0.04) | 4 × 10<sup>5</sup> | 27.21 (0.13) | 23.81 (0.14) |
|  | 3.22 × 10<sup>4</sup> | 31.31 (0.05) | 26.56 (0.12) | 4 × 10<sup>4</sup> | 33.92 (0.39) | 27.82 (0.17) |
|  | 3.22 × 10<sup>3</sup> | NA | 30.89 (0.40) | 4 × 10<sup>3</sup> | NA | 31.79 (0.64) |
|  | 3.22 × 10<sup>2</sup> | NA | NA | 4 × 10<sup>2</sup> | NA | NA |
|  | 3.22 × 10<sup>1</sup> | NA | NA | 4 × 10<sup>1</sup> | NA | NA |
|  | 0 to 4 | NA | NA | 0 to 4 | NA | NA |
| Efficiency<sup>c</sup> |  | 94.50 | 105.2% |  | 83.10 | 110.8% |
| R<sup>2</sup> |  | 0.994 | 0.987 |  | 0.934 | 0.98 |
| Slope |  | -3.461 | -3.202 |  | -3.808 | -3.088 |
| P135<sub>2d</sub> | 5.80 × 10<sup>7</sup> | 20.52 (0.12) | 17.1 (0.13) | 2.04 × 10<sup>7</sup> | 20.97 (0.16) | 18.28 (0.14) |
|  | 5.80 × 10<sup>6</sup> | 23.25 (0.06) | 19.93 (0.11) | 2.04 × 10<sup>6</sup> | 23.78 (0.17) | 20.99 (0.18) |
|  | 5.80 × 10<sup>5</sup> | 26.13 (0.01) | 22.72 (0.28) | 2.04 × 10<sup>5</sup> | 27.32 (0.14) | 24.68 (0.32) |
|  | 5.80 × 10<sup>4</sup> | 30.30 (0.21) | 26.79 (0.16) | 2.04 × 10<sup>4</sup> | 31.78 (0.50) | 27.51 (0.13) |
|  | 5.80 × 10<sup>3</sup> | NA | 28.95 (0.08) | 2.04 × 10<sup>3</sup> | NA | 31.09 (0.23) |
|  | 5.80 × 10<sup>2</sup> | NA | 33.28 (0.56) | 2.04 × 10<sup>2</sup> | NA | NA |
|  | 5.80 × 10<sup>1</sup> | NA | NA | 2.04 × 10<sup>1</sup> | NA | NA |
|  | 0 to 4 | NA | NA | 0 to 3 | NA | NA |
| Efficiency<sup>c</sup> |  | 104.40 | 107.6% |  | 89.60 | 104.7% |
| R<sup>2</sup> |  | 0.987 | 0.99 |  | 0.980 | 0.994 |
| Slope |  | -3.221 | -3.153 |  | -3.598 | -3.214 |

<sup>a</sup> Concentration in samples before DNA extraction. Seed extract samples were spiked with serial dilutions of each strain separately. DNAs were extracted from 1 mL of seed sample, resuspended in 30 µL of elution buffer and tested in triplicates with 1 µL per reaction.
<sup>b</sup> NA = Not Amplified
<sup>c</sup> Standard curve and qPCR parameters were determined in seed extract matrix.

Four seed lots (I to IV) were analyzed with the Multiplex qPCR test after DNA extraction on seed extracts. All seed lots were naturally contaminated (Table 7). Based on the number of seeds per subsample, the number of subsamples, and the number of positive subsamples, contamination rates of seed lots I to IV were estimated at 1.24%, 1.24%, 0.9% and 2.4%, respectively, according to Maury’s formula for subsamples of the same size (Maury et al. 1985), and to Swaroop’s tables for subsamples with decreasing sizes (Swaroop, 1951). Seed lots I and II were contaminated at the same rate (1.2%). For each lot, bacterial concentrations were similar in the same subsamples for *hopZ5* and *sylC* PCR tests. This strongly suggested an infection with a VCZ strain with a wide host range. Seed lot III was contaminated at 0.9% probably with a clade 2d strain because population sizes estimated with both tests (2d and *sylC* PCR tests) were congruent in the sub-samples. Seed lot IV corresponded to a disinfected seed lot that was resistant to disinfection. Because the number of genome copies estimated for *sylC* in subsamples could correspond to the sum of the genome copies estimated for *avrRpt2* and *hopZ5* markers, i.e the sum of 0.91 × 10^6^ CFU per 500 seeds (number of genomes having avrRpt2 marker) and 0.2 × 10^6^ CFU per 500 seed (number of genomes having *hopZ5* marker) was 1.11 × 10^6^ CFU which is close to the 1.17 × 10^6^ CFU per 500 seeds estimated for the marker *sylC*. This lot was then probably infected with two strains, one having a narrow host range that was more abundant than a second strain having a wider host range.

**Table 7.** Results of the analysis of naturally contaminated seed lots with the Multiplex qPCR test

| Seed lot<sup>a</sup> | Number of seeds per subsample | Mean log genome copies per seed sample<sup>b</sup> (SEM) |  |  |  |
| --- | --- | --- | --- | --- | --- |
|  |  | 4Ba | Z5 | Rpt2 | 2d |
| I | 100 | 5.85 (0.05) | 5.66 (0.03) | NA | NA |
|  | 100 | 5.34 (0.38) | 5.63 (0.02) | NA | NA |
|  | 100 | 5.58 (0.33) | 5.76 (0.01) | NA | NA |
|  | 100 | 5.85 (0.23) | 5.77 (0.03) | NA | NA |
|  | 100 | 5.30 (0.41) | 5.52 (0.05) | NA | NA |
|  | 100 | NA<sup>c</sup> | NA | NA | NA |
|  | 100 | NA | NA | NA | NA |
| II | 100 | 5.82 (0.21) | 5.83 (0.00) | NA | NA |
|  | 100 | 5.77 (0.39) | 5.85 (0.01) | NA | NA |
|  | 100 | 5.99 (0.25) | 5.99 (0.02) | NA | NA |
|  | 100 | 5.78 (0.18) | 5.90 (0.04) | NA | NA |
|  | 100 | 5.58 (0.45) | 5.81 (0.02) | NA | NA |
|  | 100 | NA | NA | NA | NA |
|  | 100 | NA | NA | NA | NA |
| III | 500 | 5.57 (0.00) | NA | NA | 5.62 (0.05) |
|  | 100 | 5.66 (0.55) | NA | NA | 5.54 (0.27) |
|  | 100 | 5.19 (0.89) | NA | NA | 6.11 (0.76) |
|  | 100 | 5.64 (0.00) | NA | NA | 5.39 (0.25) |
|  | 100 | NA | NA | NA | NA |
|  | 100 | NA | NA | NA | NA |
| IV | 500 | 6.07 (0.18) | 4.32 (0.02) | 5.96 (0.64) | NA |
|  | 100 | 5.17 (0.35) | 2.79 (0.60) | 4.88 (0.35) | NA |
|  | 100 | 5.16 (0.36) | 2.57 (0.19) | 4.77 (0.36) | NA |
|  | 100 | 5.50 (0.27) | 3.52 (0.40) | 5.07 (0.34) | NA |
|  | 100 | 5.01 (0.34) | 3.50 (0.36) | 4.54 (0.36) | NA |
|  | 100 | 5.48 (0.31) | 3.56 (0.41) | 5.34 (0.51) | NA |
|  | 10 | NA | NA | NA | NA |
|  | 10 | NA | NA | NA | NA |
|  | 10 | NA | NA | NA | NA |
|  | 10 | NA | NA | NA | NA |
|  | 10 | NA | NA | NA | NA |

<sup>a</sup> Seed lots I to III were produced in France in 2019 and IV in China in 2020.
<sup>b</sup> Number of genome copies per sample of 100 or 500 seeds was the number genome copies in 1 µL of PCR reaction multiplied by the total volume of extracted DNA and by the total volume of seed soaking. Samples were amplified in two different experiments and two different equations were used to calculate the number of genome copies per reaction.
<sup>c</sup> NA = Not Amplified

## Discussion

Strains of *Pseudomonas syringae* are ubiquitous plant epiphytes and some are plant pathogens. They are diverse and, according to the weapons they dispose, such as T3SS and effectors, toxins, exopolysaccharides, cell wall-degrading enzymes and plant hormones (Xin et al 2018), they do not represent the same risk for a crop. The aim of the study was to provide tools to characterize, identify, detect and quantify strains belonging to the *P. syringae* species complex that affect zucchini seed production: (i) PCR tests were designed to extend the previously proposed MLSA scheme to seven genes to characterize isolates from epidemics and establish their phylogenetic relationships and their taxonomy (Lacault et al. 2020). (ii) Based on a set of genome sequences, PCR tests have been developed to determine, without the need for sequencing, to which phylogenetic group the strains belong. These user-friendly tests are dedicated to large numbers of samples for epidemiological monitoring. (iii) In order to improve seed testing, the qPCR test based on *sylC* was multiplexed with other markers in order to identified the strains according to their pathogenicity.

The new MLSA scheme based on seven genes allowed the characterization of the whole collection of VCZ strains and additional strains isolated from zucchini seeds and plants from seed production fields. Two VCZ strains were identified in the cluster 2ba-C, indicating that VCZ strains are present in five clusters among the six that were previously described for strains pathogenic on cucurbits (Lacault et al. 2020, Newberry et al. 2019). Clustering of strains in homogeneous lineages suggests they could be epidemic clones responsible for damages on zucchini seedlings (Lacault et al. 2020) and for some of them also on other cucurbits (Lacault et al. 2020; Newberry et al. 2019). MLSA revealed that strains belonging to clades 2b and 2d did not form clusters. The clades-2d strains can cause symptoms on cucurbits, but were less aggressive on zucchini plantlets than VCZ control strains. Strains P129_2d_ and P135_2d_ were discarded from our previous study (Lacault et al 2020) because they were not considered as VCZ strains. Indeed, they did not cause plantlet deformation and stunting as VCZ control did (Supplementary Figure 1A). However, they were able to cause water-soaked and necrotic lesions on leaves of various cucurbits after foliar spray inoculation, including several squashes, melons, cucumber and a watermelon, as previously described (Lacault et al 2020) (Supplementary Figure 1B). Weak pathogenicity on squashes was previously reported for clade-2d strains (Newberry et al. 2016). Genome analysis of clade-2d strains revealed small T3E repertoires similarly to PG2 genome sequences in general (Dillon et al. 2019a). They had fewer T3Es in common with VCZ-clusters strains than the clade-2b strain P108_2b_. Strains of clades 2b and 2d lacked *hopA1* which is shared by all VCZ strains (Lacault et al. 2020) and was reported to be present in all strains responsible for epidemics on cucurbits in the USA (Newberry et al. 2019). Being never organized in genetic clusters, clades-2b and −2d strains did not seem to be epidemic clones, even if they can cause limited symptoms following inoculation. Improving the number of these types of strains in collection is necessary to precise their role in a potential zucchini disease.

In order to detect VCZ and related strains, we used genes encoding pathogenicity factors to improve the qPCR test based on *sylC*. This gene encodes a NRPS (nonribosomal peptide synthetase) module involved in the biosynthesis of syringolin A, a toxin secreted only by strains belonging to phylogroup 2 (Dillon et al. 2019b). The *sylC* test (4Ba) detects all VCZ strains clustered in the clades 2a and 2ba and also some other strains within PG2, in particular all clade 2d strains and a large part of clade 2b strains (Table 1 and Supplementary Table 2). The test based on *sylC* was multiplexed with qPCR tests targeting *avrRpt2* and *hopZ5*. The gene encoding the T3E AvrRpt2 is present in all VCZ strains that have a narrow host range on cucurbits (Lacault et al. 2020; Djitro et al. 2022a) but also in the pathovar *tomato* (Innes et al. 1993) and was found in some strains of pathovars *maculicola, spinaceae* and *tagetis* that were not known to have it. Indeed, *avrRpt2* was not included in the large analysis of the distribution of virulence encoding genes including 53 T3Es in *Pseudomonas syringae* made by Sarkar et al. (2006). A BLASTn confirmed that those strains have the complete gene sequence in their genome. As none of these strains, including those of the pathovar *tomato,* gave a positive signal *in silico* with the qPCR test based on *sylC* and neither *in vitro* for the strain of pathovar *tomato*, their reaction pattern using our multiplex test would be different to that of VCZ strains. The gene encoding the T3E *hopZ5* is present in VCZ strains having a wide cucurbit host range and in the pathovar *actinidiae,* in which this T3E was described (Jarayaraman et al. 2017). The pathovar *actinidiae* was not detected with the *sylC* qPCR, and hence gave a different pattern than VCZ strains with our multiplex test. We tried to improve the information provided by the test with the addition of qPCR tests for clades 2b and 2d. However, only the test for clade 2d showed an efficiency consistent with the multiplex. We included this clade 2d qPCR test in the multiplex to identify cucurbit-associated strains to gain insight on their occurrence and further verify if they represent a low risk for zucchini seed sector.

Specificity of the multiplex qPCR test was based on the combined results of different tests that were not individually specific, except the qPCR test designed to detect clade 2d strains. To be perfectly reliable, this multiplex qPCR test should be performed on isolated clones, because on crude seed extracts, it cannot be excluded that positive signals could arise from a mixture of strains. Such a warning was previously stated for a multiplex PCR test, based on the simultaneous detection of two genes encoding T3Es, dedicated to the identification of common bacterial blight agents (Boureau *et al*. 2013). However, the quantification of each marker that is possible with qPCR could help to solve this question and allow to identify if marker-corresponding population sizes are compatible with single or mix contaminations. Furthermore, subsampling which is performed to determine seed lot contamination rates (ISTA, 2015) could bring complementary arguments about single or mix contaminations according to their distributions simultaneously in the same subsamples or separately in different subsamples as illustrated in Table 7.

More generally, detection tests based on T3Es or other virulence factors present de facto a risk of non-specificity, as that these pathogenicity factors are easily transferable among strains, the repertoire of which evolved rapidly (Baltrus *et al*. 2011; Dillon et al. 2019a). Recombination and horizontal gene transfers were highlighted by several comparative genomic studies within the *P. syringae* species complex (Baltrus et. al 2017; Hulin et. al 2018; Dillon et al. 2019b; Newberry et al. 2019). However, targeting genes involved in pathogenicity could give clues to discriminate strains according to their interactions with plants, as was shown for *P. syringae* (Sarkar et al. 2006) and *Xanthomonas* (Hajri et al. 2009), but not for *Ralstonia solanacearum* (Peeters et al. 2013). For example, the test based on the gene encoding SyrD, which is involved in the secretion of syringomycin and syringopeptin, is used to discriminate the strains of the pathovar *syringae* from those of pv. *morsprunorum* on cherry tree or from those of pv. *pisi* on pea (Bulthreys and Gheysen 1999). Another example is the harpin HrpZ, whose encoding gene is the preferred target for the detection of the pathovar *tomato*. Primers are designed in specific regions within this gene which is distributed among many pathovars. This gene is still in use to develop new molecular diagnosis tools (Zaccardelli et al. 2005, Chen et al. 2020, Chai et al. 2020).

The protocol we used to test seeds was was adapted to discard PCR inhibitors present in zucchini seed extracts. The LOD of the multiplex qPCR test for spiked seed samples was as low as 2.5 × 10^3^ CFU.mL^-1^ corresponding to 7.5 × 10^3^ CFU.g^-1^ of seeds. DNA extraction protocol included a first step of centrifugation followed by several purification steps and elution in a smaller volume than the initial volume allowing to concentrate the samples (33-fold enrichment) without concentrating PCR inhibitors, which is important for PCR because sensitivity of the method is limited by the small size of the analyzed volume. Efficiencies of qPCR tests were as good on spiked seed extracts after DNA extraction as on bacterial suspensions. The adaptation of the DNA extraction protocol is crucial to the success of the test, because the presence of polymerase inhibitors from plant material can have negative repercussions on assay performance and not all protocols are adapted to all matrix (Lau and Botella, 2017). For example, for the same DNA extraction protocol, the efficiency of detection of *Xylella fastidiosa in planta* could vary according to plant matrix probably due to abundant inhibitors such as polyphenols and polysaccharides. The negative impact may decrease by 10 to 100 times the LOD of the test in plant material in comparison with bacterial suspensions (Dupas et al. 2019).

The multiplex qPCR test developed in this work could be useful for epidemiological purposes to improve our knowledge on VCZ and would constitute an informative complement for the seed sector to the qPCR test in-use that is based on *sylC*. It was possible in one step to identify different kind of strains, to quantify them and, by the means of subsampling to determine the contamination rates of the seed lots. Compared to pre-existing methods (qPCR based on *sylC*, strain isolation in positive samples and MLSA characterization) this method is particularly fast and cost-effective. It could be useful for epidemiological studies aiming at identifying the sources of inoculum in parental seed lots or in the environment of the seed production fields. The multiplex qPCR test will be very useful for seed testing because it provides information on the risk associated with contaminations present in the seed lots. Depending on whether the seeds are contaminated by a strain with *avrRpt2* or *hopZ5*, the risk of damages for surrounding crops could be limited to *Curcurbita* sp or be extended to other cucurbits such as melons, cucumber and watermelon, especially since epidemics involving various cucurbits crops have reported in USA for closely related strains (Newberry et al. 2016 and 2019). The subsampling of seed lots allows detection and quantification of mixed infections. The detection limit of this method is to 5 × 10^5^ CFU per sample of 500 seeds (for a 1,000-seed weight of 145 g). It would be interesting to determine the epidemiological risk associated with this detection threshold, for example by the survey of fields sown with contaminated seed lots, and would inform on the need of improving the sensitivity of this detection tool for seed testing.

