## Supplementary Figure 1 for "Development of tools to detect and identify strains belonging to the *Pseudomonas syringae* species complex responsible for vein clearing of zucchini"

### Slide 1
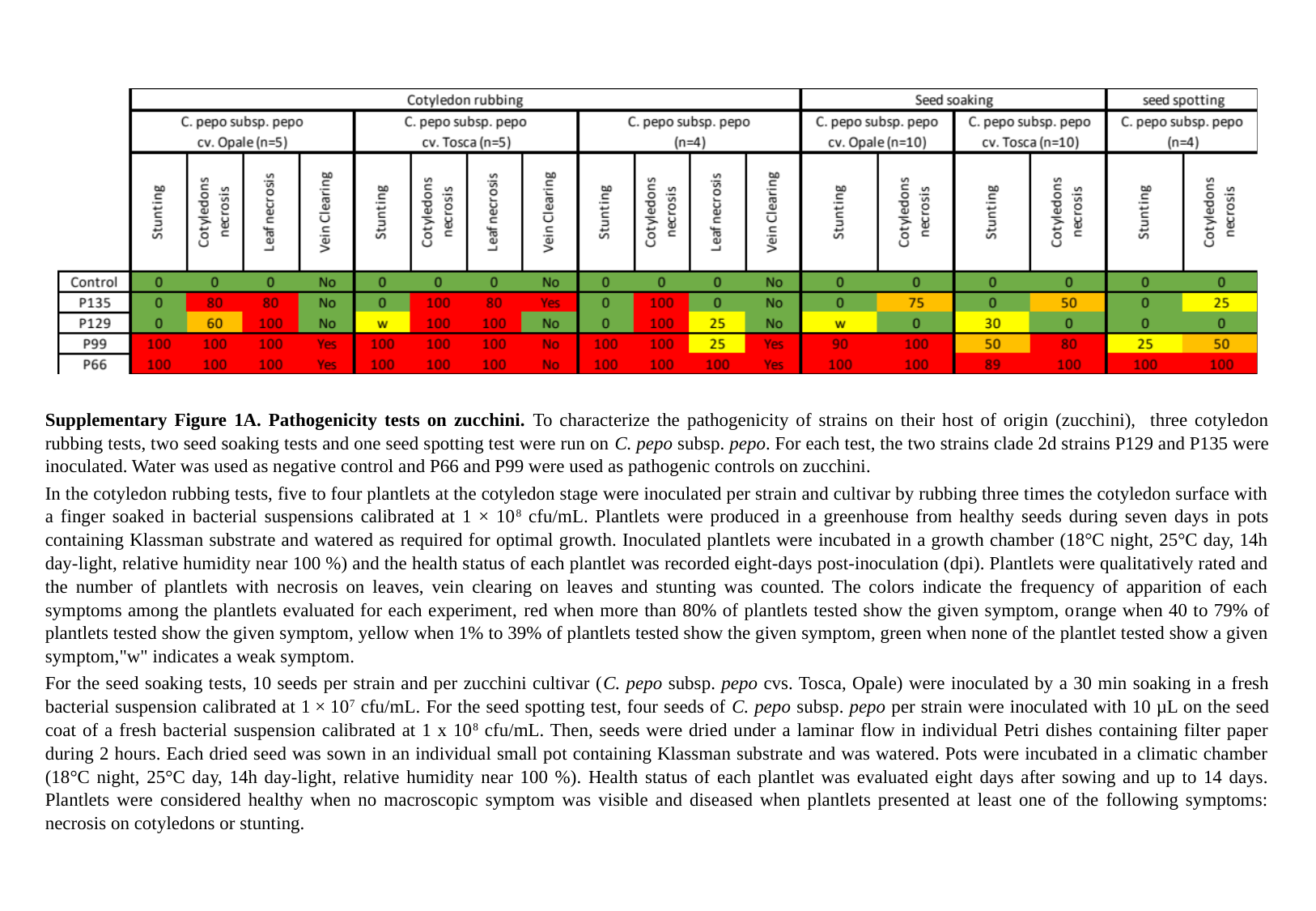

Supplementary Figure 1A. Pathogenicity tests on zucchini. To characterize the pathogenicity of strains on their host of origin (zucchini), three cotyledon rubbing tests, two seed soaking tests and one seed spotting test were run on C. pepo subsp. pepo. For each test, the two strains clade 2d strains P129 and P135 were inoculated. Water was used as negative control and P66 and P99 were used as pathogenic controls on zucchini.
In the cotyledon rubbing tests, five to four plantlets at the cotyledon stage were inoculated per strain and cultivar by rubbing three times the cotyledon surface with a finger soaked in bacterial suspensions calibrated at 1 × 10^8^ cfu/mL. Plantlets were produced in a greenhouse from healthy seeds during seven days in pots containing Klassman substrate and watered as required for optimal growth. Inoculated plantlets were incubated in a growth chamber (18°C night, 25°C day, 14h day-light, relative humidity near 100 %) and the health status of each plantlet was recorded eight-days post-inoculation (dpi). Plantlets were qualitatively rated and the number of plantlets with necrosis on leaves, vein clearing on leaves and stunting was counted. The colors indicate the frequency of apparition of each symptoms among the plantlets evaluated for each experiment, red when more than 80% of plantlets tested show the given symptom, orange when 40 to 79% of plantlets tested show the given symptom, yellow when 1% to 39% of plantlets tested show the given symptom, green when none of the plantlet tested show a given symptom,"w" indicates a weak symptom.
For the seed soaking tests, 10 seeds per strain and per zucchini cultivar (C. pepo subsp. pepo cvs. Tosca, Opale) were inoculated by a 30 min soaking in a fresh bacterial suspension calibrated at 1 × 10^7^ cfu/mL. For the seed spotting test, four seeds of C. pepo subsp. pepo per strain were inoculated with 10 µL on the seed coat of a fresh bacterial suspension calibrated at 1 x 10^8^ cfu/mL. Then, seeds were dried under a laminar flow in individual Petri dishes containing filter paper during 2 hours. Each dried seed was sown in an individual small pot containing Klassman substrate and was watered. Pots were incubated in a climatic chamber (18°C night, 25°C day, 14h day-light, relative humidity near 100 %). Health status of each plantlet was evaluated eight days after sowing and up to 14 days. Plantlets were considered healthy when no macroscopic symptom was visible and diseased when plantlets presented at least one of the following symptoms: necrosis on cotyledons or stunting.

### Slide 2
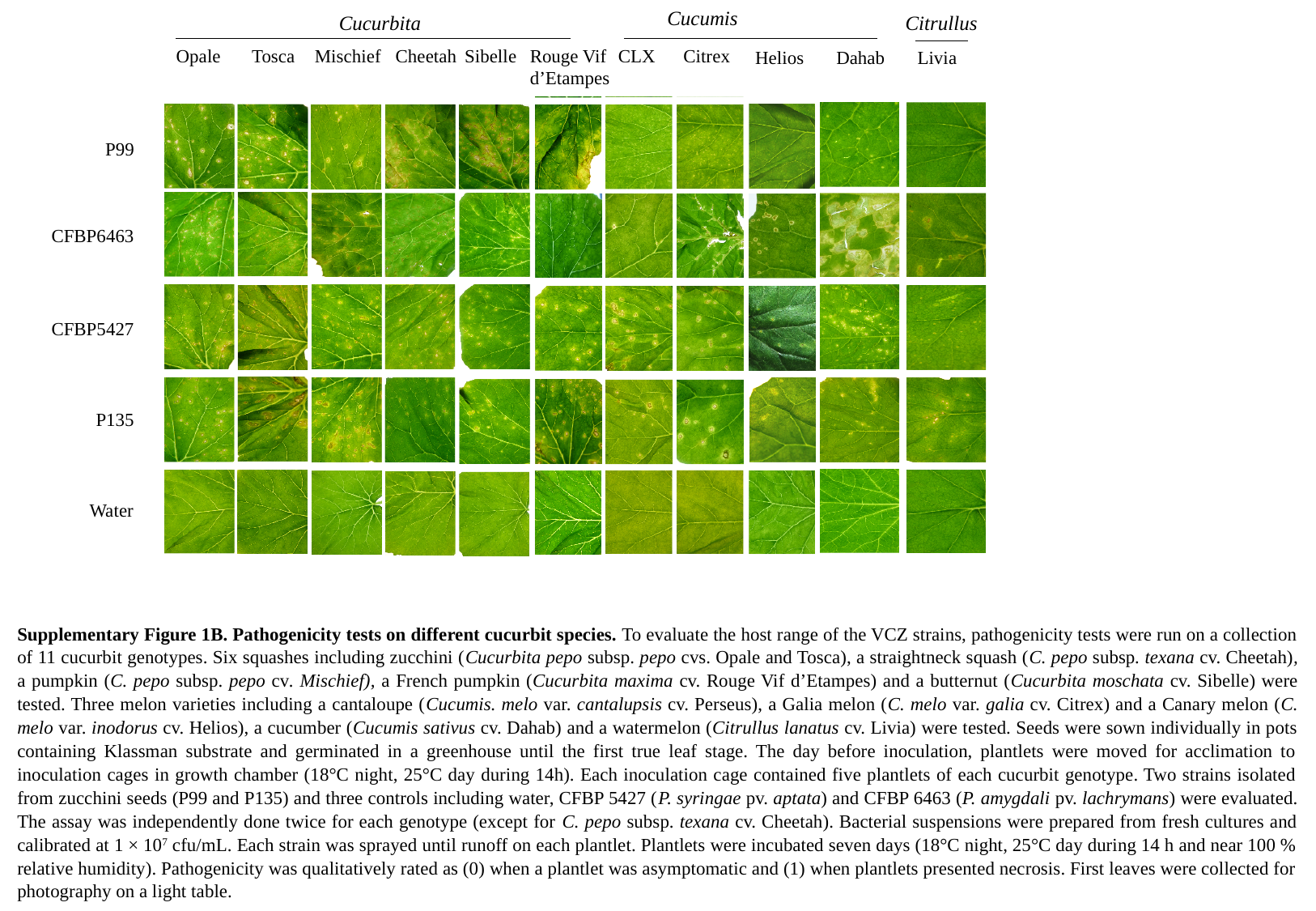

Cucumis
Cucurbita
Citrullus
Opale
Tosca
Mischief
Cheetah
Sibelle
Rouge Vif
d’Etampes
Citrex
CLX
Helios
Dahab
Livia
P99
CFBP6463
CFBP5427
P135
Water
Supplementary Figure 1B. Pathogenicity tests on different cucurbit species. To evaluate the host range of the VCZ strains, pathogenicity tests were run on a collection of 11 cucurbit genotypes. Six squashes including zucchini (Cucurbita pepo subsp. pepo cvs. Opale and Tosca), a straightneck squash (C. pepo subsp. texana cv. Cheetah), a pumpkin (C. pepo subsp. pepo cv. Mischief), a French pumpkin (Cucurbita maxima cv. Rouge Vif d’Etampes) and a butternut (Cucurbita moschata cv. Sibelle) were tested. Three melon varieties including a cantaloupe (Cucumis. melo var. cantalupsis cv. Perseus), a Galia melon (C. melo var. galia cv. Citrex) and a Canary melon (C. melo var. inodorus cv. Helios), a cucumber (Cucumis sativus cv. Dahab) and a watermelon (Citrullus lanatus cv. Livia) were tested. Seeds were sown individually in pots containing Klassman substrate and germinated in a greenhouse until the first true leaf stage. The day before inoculation, plantlets were moved for acclimation to inoculation cages in growth chamber (18°C night, 25°C day during 14h). Each inoculation cage contained five plantlets of each cucurbit genotype. Two strains isolated from zucchini seeds (P99 and P135) and three controls including water, CFBP 5427 (P. syringae pv. aptata) and CFBP 6463 (P. amygdali pv. lachrymans) were evaluated. The assay was independently done twice for each genotype (except for C. pepo subsp. texana cv. Cheetah). Bacterial suspensions were prepared from fresh cultures and calibrated at 1 × 10^7^ cfu/mL. Each strain was sprayed until runoff on each plantlet. Plantlets were incubated seven days (18°C night, 25°C day during 14 h and near 100 % relative humidity). Pathogenicity was qualitatively rated as (0) when a plantlet was asymptomatic and (1) when plantlets presented necrosis. First leaves were collected for photography on a light table.
